# Microbial Signatures of Oral Dysbiosis, Periodontitis and Edentulism Revealed by Gene Meter Methodology

**DOI:** 10.1101/070367

**Authors:** M. Colby Hunter, Alex E. Pozhitkov, Peter A. Noble

## Abstract

Conceptual models suggest certain microorganisms (e.g., the red complex) are indicative of a specific disease state (e.g., periodontitis); however, recent studies have questioned the validity of these models. Here, the abundances of 500+ microbial species were determined in 16 patients with clinical signs of one of the following oral conditions: periodontitis, established caries, edentulism, and oral health. Our goal was to determine if the abundances of certain microorganisms reflect dysbiosis or a specific clinical condition that could be used as a signature for dental research. Microbial abundances were determined by the analysis of 138,718 calibrated probes using Gene Meter methodology. Each 16S rRNA gene was targeted by an average of 194 unique probes (*n*=25 nt). The calibration involved diluting pooled gene target samples, hybridizing each dilution to a DNA microarray, and fitting the probe intensities to adsorption models. The fit of the model to the experimental data was used to assess individual and aggregate probe behavior; good fits (R^2^>0.90) were retained for back-calculating microbial abundances from patient samples. The abundance of a gene was determined from the median of all calibrated individual probes or from the calibrated abundance of all aggregated probes. With the exception of genes with low abundances (< 2 arbitrary units), the abundances determined by the different calibrations were highly correlated (*r* ∼1.0). Seventeen genera were classified as signatures of dysbiosis because they had significantly higher abundances in patients with periodontitis and edentulism when contrasted with health. Similarly, 13 genera were classified as signatures of periodontitis, and 14 genera were classified as signatures of edentulism. The signatures could be used, individually or in combination, to assess the clinical status of a patient (e.g., evaluating treatments such as antibiotic therapies). Comparisons of the same patient samples revealed high false negatives (45%) for next-generation-sequencing results and low false positives (7%) for Gene Meter results.

## INTRODUCTION

Periodontitis is an inflammatory disease associated with the tissues that support the teeth. The disease causes the progressive loss of bone that could result in tooth loss (1). A small assemblage of bacterial species was once thought to be the cause of periodontal disease. This assemblage consisted of *Porphyromonas gingivalis, Tannerella forsythia,* and *Treponema denticola,* and was ominously dubbed the “red complex” (2,3,4). The prevailing idea at the time was that the red complex changed the signaling pathways of the host. This change made the host vulnerable and led to a change in the relative abundance of microorganisms thus causing the disease (5,6). However, members of the red complex have been discovered in people who do not have periodontal disease (8,7,9). Therefore, members of the red complex could be classified as pathogens or harmless commensals. This fact, along with the finding that the oral microbiome of humans is much more diverse than previously thought (10,11), has prompted a paradigm shift in the understanding of the origins of periodontal disease. This new paradigm states that periodontal disease is not caused by the presence of specific bacteria, but by changes in the population levels of species in the oral microbiome (12). What begins as a symbiosis between host and microbes becomes a dysbiosis as the microorganisms transcend beyond the host-imposed boundaries. The premise of our study is that a change in microbial populations can be determined by the relative abundances of individual species in patients with different clinical conditions.

Oligonucleotide microarray technology has been used to profile microbial communities for quite some time (13,14,15,16,17). A new version of the methodology, the Gene Meter, was recently introduced (although not explicitly named “Gene Meter” in refs. 18,19). Unlike conventional DNA microarrays, a calibration procedure is conducted prior to applying samples onto the microarrays (20,21). The calibration procedure establishes the microarray probe responses to increasing target concentrations (i.e., a dilution series). Individual probes are calibrated or, alternatively, groups of probes collectively specific to a target (i.e., aggregates) are calibrated. Both approaches of calibration were compared in this study.

Next generation sequencing (NGS) has also been used to profile oral microbial communities (7,9,22,23,24,25,26,27). This approach involves the massive sequencing of thousands to millions of DNA strands in a single run (28). While microbial abundances may be quantified by NGS, it remains to be determined if the quantifications truly represent the quantities of the genes in a biological sample. Only a limited number of studies have rigorously challenged their quantitative capabilities. In one study, a complex microbial mixture was spiked with 5 fungal targets including *L. proxima* and *M. noctilunum* (29). It was shown that depending on the target, the coefficient of determination for the relationship between actual quantities of the target and its sequencing counts was very low. In another recent study, NGS is shown to be a semi-quantitative approach for examining fungal populations (30). The results of the study showed that 454 pyrosequencing counts are, at best, moderately related with spore concentration measurements by qPCR (R^2^ ∼0.5). An RNA-seq spike-in study has also been conducted (31). Ninety-six RNA sequences of concentrations varying 6 orders of magnitude were mixed and sequenced and the concentrations were correlated to the number of reads. Although the study design did not investigate the response of an individual target to concentration, the results suggest that the RNA-seq sensitivity can vary up to 10-fold depending on the gene target (31). The uncertainty associated with quantification by sequencing raises questions about the validity of such quantifications and warrants a well-controlled calibration study, which was done here.

The objectives of the study are three-fold: (i) to demonstrate the utility of the Gene Meter approach to precisely determine 16S rRNA gene abundances using two calibration approaches, (ii) to determine if certain microorganisms have an abundance signature that can be employed to identify specific or general oral health conditions, and (iii) to compare and contrast two different technologies (Gene Meter and DNA sequencing) using the same 16 patient samples.

## MATERIALS AND METHODS

The DNA sequencing, the study design, and sample collection methods have been previously published (32). The same amplified DNA used for DNA sequencing was also analyzed by the Gene Meter approach. Splitting the samples in this way enabled direct comparison of results from two distinctly different molecular methods. To familiarize readers with the previous study, we have provided a ßbrief overview of patient recruitment, enrollment, and exclusion criteria, sample collection, DNA extraction and amplification, and DNA sequencing analyses, below.

### Patient recruitment, enrollment, and exclusion criteria

The adult patients were recruited at the University of Washington, Seattle, USA, and the University of Düsseldorf, Germany and enrolled if they had one of the following clinical conditions: severe periodontitis, caries, edentulism, and oral health. A periodontitis case was defined as having at least two interproximal sites at different teeth with clinical attachment loss (CAL) of 6 mm or greater and at least one interproximal site with probing depth (PD) of 5 mm or greater (33) and a minimum of 20 permanent teeth, not including third molars. Patients were excluded from the periodontitis group if they had any established caries lesion or wore a removable partial denture. A caries case was defined as having the following number of teeth with established caries lesions: 6 or more teeth in subjects 20 to 34 years of age; 4 or more teeth in subjects 35 to 49 years of age; and 3 or more teeth in subjects 50 years of age and older. Established caries was defined as a class 4 lesion according to the International Caries Detection and Assessment System. The number of teeth with caries lesion in caries cases was greater than one standard deviation above the mean of caries extent in respective age group the U.S.A (34). Exclusion criteria for a caries case were interproximal sites with CAL of 4 mm or greater or PD of 5 mm or greater (33). An edentulous case had to be completely edentulous in both jaws and their teeth had to be extracted more than one year before the enrollment in the study. A healthy case was defined as having 28 teeth, not counting third molars, or 24 or more teeth, not counting third molars if premolars had been extracted for orthodontic reasons or were congenitally missing with no signs of oral disease. Exclusion criteria for a healthy case included: smoking, loss of permanent teeth due to caries or periodontal disease, any interproximal sites with CAL of four or greater or PD of 5 mm or greater, or any established caries lesions. Exclusion criteria for all groups included: oral mucosal lesions, systemic diseases, and use of antibiotics or local antiseptics within 3 months prior to the study.

### Sample collection, DNA extraction and PCR amplification

For all but the edentulous patients, supra- and subgingival plaque was collected from sites with the deepest probing depth in each sextant. One sterile paper point per site was inserted into the deepest aspect of the periodontal pocket or gingival sulcus. Biofilm from oral mucosae was collected by swiping a sterile cotton swab over the epithelial surfaces of the lip, left and right buccal mucosae, palate, and dorsum of the tongue (35). Samples were stored at -80°C.

Microbial DNA was isolated from cells by physical and chemical disruption using zirconia/silica beads and phenol-chloroform extraction in a FastPrep-24 bead beater (36). Prokaryotic 16S rRNA genes were amplified using universal primers (27F and 1392R) using the GemTaq kit from MGQuest (Cat# EP012). The PCR program involved a pre-amplification step of 10 cycles with an annealing temperature of 56^o^C followed by 20 amplification cycles with annealing temperature 58^o^C. In each cycle, elongation time was 1 min 10s, at 72^o^C. PCR was finalized by extended elongation for 5 min. PCR products were purified with DNA Clean & Concentrator columns (Zymo Research, USA) and quantified using the NanoDrop (Agilent, USA). Equal quantities of PCR product derived from swab and paper point samples were pooled together for each patient. For edentulous patients, there were no paper point samples. Each purified PCR product was sequenced on a Roche 454 Jr. instrument as previously described (32).

### DNA sequencing analysis

The obtained sequences were uploaded to the MG-RAST web server (37). The MG-RAST pipeline assessed the quality of sequences, removed short sequences (multiplication of standard deviation of length cutoff of 2.0) and removed sequences with ambiguous bp (non-ACGT; maximum allowed number of ambiguous base pair was set to 5). The pipeline annotated the sequences and allowed the integration of the data with previous metagenomic and genomic samples. The RDP database was used as an annotation source, with minimum sequence identity of 97%, maximum e-value cutoff of 10-5, and minimum sequence length of 100 nt.

### Microarray probe design

Microarray probes were obtained by tiling along 16S rRNA sequences of 597 oral bacteria (Table S1) using a program written for this purpose. Redundant probes were removed. Each probe was 25 nt in length. The 16S rRNA gene sequences used to design the probes ranged from 424 to 13,214 nt in length with an average of 1,492 nt. Three of the 16S rRNA sequences also included 5S and 23S rRNA genes. The microarrays were created by NimbleGen (now Roche Inc.).

### Sample labeling

PCR products (above) were purified using a “DNA Clean & Concentrator” kit (Zymo Research, USA), dried under a flow of dry nitrogen and labeled using ULYSIS direct chemical labeling kit. The attached dye was Alexa Fluor 546 (i.e., spectral analog of Cy3).

### DNA microarray calibration

Probes on the microarrays were calibrated using a dilution series of labeled pooled PCR products of all samples. The dilution series for the NimbleGen array was created using the following quantities of labeled PCR amplicons: 11.99, 7.72, 3.09, 1.54, 0.77, and 0.39 µg in 12.5 µl. Signal intensities were collected and stored in a database. Two independent calibration procedures were conducted i.e., calibration of individual probes and calibration of probe aggregates. An automatic fitting procedure determined the best fit curve – the calibration curve – (e.g., Langmuir, Linear or Freundlich) as well as the curve parameters from the dilutions and signal intensities. For the individual probes calibration, signal intensity was that of the individual probe. For the probe aggregate calibration, signal intensity was a sum of signal intensities of the probes specific to the corresponding targets. To calculate the abundances of the targets, the calibration curve equations were inverted such that from the signal intensity a dilution factor could be obtained. The dilution factor is proportional to the concentration of the target; hence, it was defined as the abundance of the target.

Details of the calibration protocols to calculate gene abundance are provided in our recent papers (18,19). The quantities of samples loaded onto the microarray of processing of the individual patients were 3.2 ± 0.25 µg.

### Bioinformatic analyses

Matching the probes to the targets (*in silico*) and determining the gene abundances using the unique calibrated probes was conducted using custom-designed programs written in C++. T-tests were analyzed and histograms were made using SAS JMP 7 and MS Excel.

## RESULTS

### Microarray probe coverage and selection

In total, 276,234 probes (25 nt) were synthesized on NimbleGen microarrays. *In silico* matches of the probes to all 597 oral bacterial sequences (used to design the probes) revealed 175,206 probes were unique to a single 16S rRNA gene target. To determine if the microarray design had sufficient coverage, we downloaded the curated core oral microbiome 16S rRNA gene sequence dataset from http://microbiome.osu.edu and matched the unique probes to the 1,045 microbial sequences (38). We found that 66,716 probes were unique to the 1,021 microorganisms in the core oral database. In other words, the unique probes on our microarray had 97.7% coverage of the sequences in the curated core oral microbiome.

We further refined the “concept of unique probes” by removing 3 nucleotides on either end (5’- and 3’-) of the probe (*in silico*) and matching the shorter probes to the 597 16S rRNA bacterial sequences. The reason for removing these nucleotides was that nucleotides at either end could potentially cross hybridize to non-specific targets in the microarray experiments (39). In this experiment, probes were considered unique if the 19 nt core matched one 16S rRNA gene target in the dataset (i.e., 597 oral bacterial sequences). Using the 19 nt core, we found 36,488 of the original 175,206 probes could be susceptible to cross-hybridization, which left 138,718 unique probes that were used in all subsequent analyses to determine gene abundance. Therefore, the entire sequence (25 nt) of the 138,718 unique probes were used in all subsequent experiments.

### Calibration of microarray probes

All probes were calibrated using a dilution series as outlined in our previous studies (18,19,20,21). The Langmuir and Freundlich isotherm equations were fitted to the data and the one with the best fit (R^2^) was retained. Examples of two calibrated probes are shown in Figure 1. Briefly, the fitting algorithm transforms the data to obtain a straight line; two parameters, *a* and *b*, are calculated. Depending on the type of the curve corresponding to the best fit, parameters *a* and *b* are utilized according to the formula of the curve. The parameters of the retained equations and fits are shown in Figure 1. Apparently, the signal intensities of the dilution data for Probe 62 were best explained by Langmuir and the dilution data for Probe 66 were best explained by Freundlich. For the Freundlich equation,

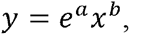

for the Langmuir equation,

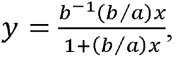

where *y* is the signal intensity, and *x* is the dilution.

These equations were used to back-calculate the gene target concentration from signal intensity values. In the case of Probe 62 at PMT=600, *a*= 0.00797, *b*=0.00034, *x*=dilution factor in the calibration series, the signal intensity (*SI*) responds as follows: *SI*= 2906.757*0.431679* *x* /(1+0.431679* *x*). Inverting this equation allows one to calculate the abundance of a gene targeted by this probe. For example, for one of the healthy patients, signal intensity was 1793.89 RFU. This yields a gene abundance of 1793.89 / ((2906.757 - 1793.89)* 0.431679)= 3.7 a.u.

**Figure 1.**
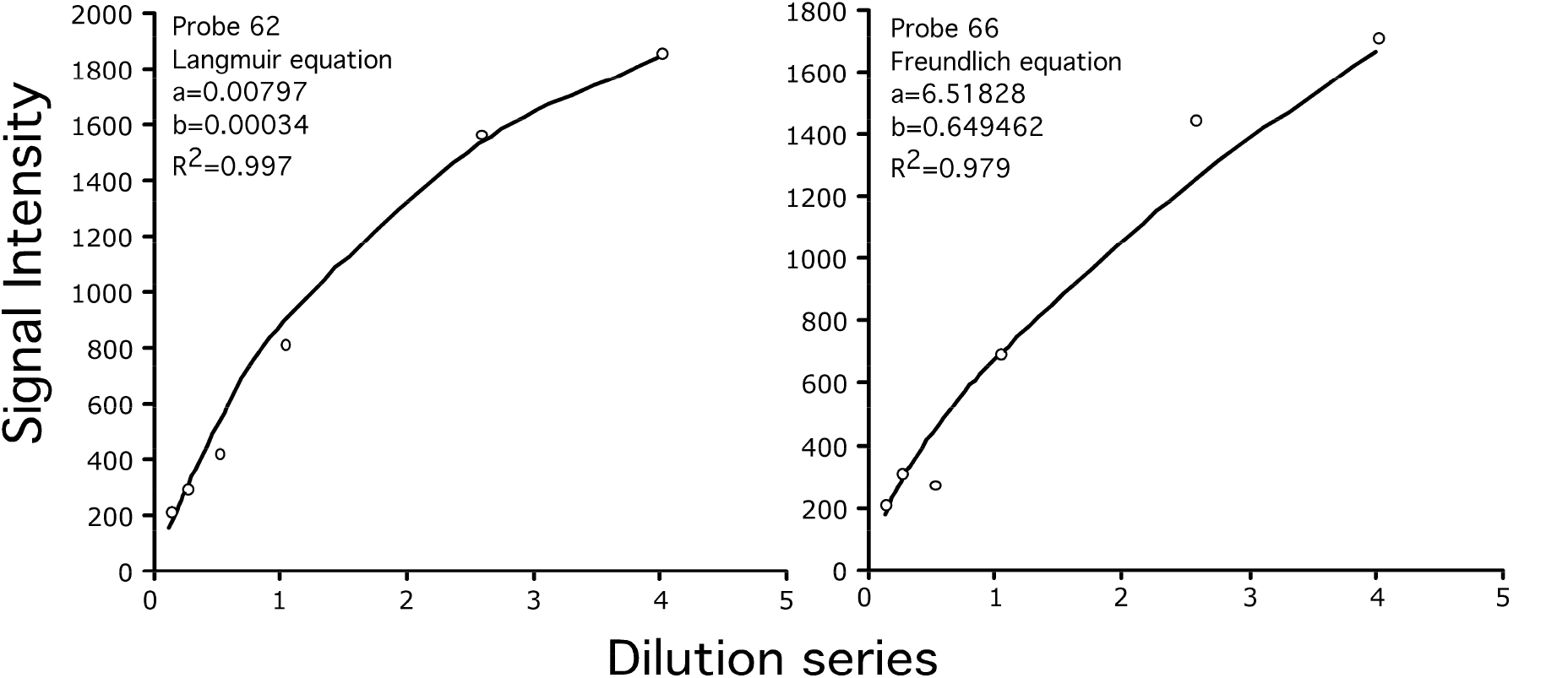
Calibration curves of two probes. Probe 62 was best calibrated using the Langmuir equation while probe 66 was best calibrated by the Freundlich equation. The equations can be used to back-calculate target concentration (dilution series) from signal intensity.

In the case of Probe 66 at PMT=600 where *a*= 6.51828, *b*=0.649462, *x*=dilution factor in the calibration series, the *SI* responds according to Freundlich equation as *SI*=exp(6.51828)\**x*^0.649462^. Inverting this equation allows one to calculate the abundance of a gene targeted by this probe. For example, for one of the healthy patients, SI= 1268.67 RFU. This yields a gene abundance of (1268.67 / exp(6.51828))^1/0.649462^=2.6 a.u.

Probe aggregates were calibrated same way as individual probes. For the probe aggregates, only linear (i.e., *y*=*ax*+*b*) and Freundlich curves were evaluated.

### Individually calibrated probes

Three PMT settings (i.e., 500, 600, 700) were used to assess the signal intensities of the hybridized duplexes because we did not know *a priori* which settings were optimal (i.e., the setting that yielded the minimum signal saturation with maximum dynamic range). The probes at each setting were independently calibrated. In total, 246,232, 246,233, and 207,006 probes were calibrated for the microarrays with the 500, 600, and 700 PMT settings, respectively. In terms of percent of the total probes, about 89% of the probes were calibrated at the 500 and 600 PMT settings and about 75% of the probes at the 700 setting. The lower percent of calibrated probes at the 700 PMT setting was due to saturation of the probes.

Median abundances ± median absolute deviation (MAD) for the 576 rRNA genes and 16 patient samples was assembled into a dataset (Table S2). Note that 21 of the original 597 gene targets were not targetted by the unique probes. The average number of calibrated probes per 16S rRNA gene was 194 ± 214 (avg. ± s.d.). The unique probes not yielding abundances for 236 any of the 16 patients were excluded. Any gene abundance that was greater than 10 a.u. was set to 10 a.u. because we did not expect the calibrations to forecast accurately beyond the dilution series used to create it. Moreover, 10.0 a.u. represents 10 times the amount of DNA hybridized to a microarray based on manufacturer’s recommendation. The average gene abundance for the 500, 600, and 700 datasets was 3.5 ± 2.0, 4.1 ± 2.1, and 3.5 ± 2.0 arbitrary units (a.u.), respectively. The lowest and highest gene abundances for all datasets were 0.3 and 10.0 a.u., respectively. The significance of these findings is that gene abundances could vary by as much as 30-fold and the averaged gene abundances were not significantly affected by PMT settings. Based on these results, all subsequent analyses of the individualized probes were performed using the 600 PMT data. The range of abundances for these data was 0.3 to 10 a.u.

### Examples: Selected microbial abundances

Statistical parameters (averages, standard deviations (s.d.), quartiles, and median) for all 576 rRNA genes in 16 patients were calculated and two examples are provided below. The histograms and whisker plots of the *Tannerella forsythia* 16S rRNA gene show the binned distribution of gene abundances in 16 patients based on 100 unique probes (Figure 2). The grey bars represent the frequencies of the binned abundances and the red bars represent the median abundances. The whisker plots show the mean, standard deviations, median, and quartiles of the abundances. The median abundance (± MAD) in the 4 patients with health was 3.0 ± 0.6 a.u., 3.0 ± 0.2 a.u., 2.2 ± 0.3 a.u., and 1.7 ± 0.3 a.u. (samples HM15, HF8, HF12 and HM10, respectively). The average gene abundance (average ± s.d.) for patients with health was 2.4 ± 0.6 a.u. The abundance of this gene in the 4 patients with edentulism, periodontitis, or caries was: 5.2 ± 1.7 a.u., 4.5 ± 1.4 a.u., and 3.5 ± 2.7 a.u., respectively. Two-tailed T-tests revealed significant differences in the average abundances for the 4 patients with edentulous and 4 patients with health (P<0.05), and the 4 patients with periodontitis and 4 patients with health (P<0.05). None of the other paired conditions for this gene were significantly different from one another (Table 1).

**Figure 2.**
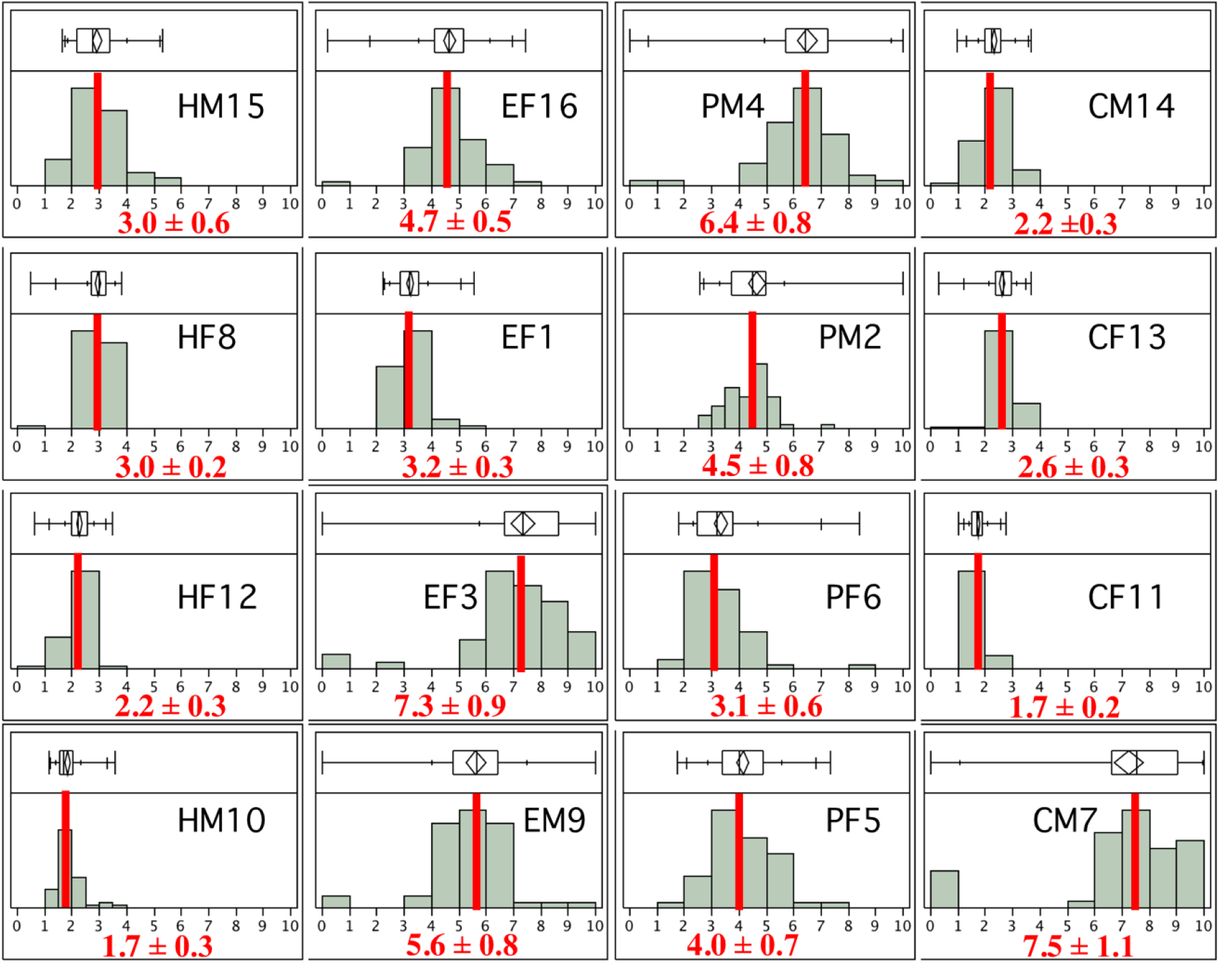
Distribution of 16S rRNA gene abundances for Tannerella forsythia (GI 289382) in 16 patients based on 50 calibrated probes. Each patient has an identifier. The first letter indicates the condition: H, health; E, edentulism; P, periodontitis; C, caries. The second letter indicates patient sex: M, male; F, female. The third number indicates patient number. The red bar indicates median value. Corresponding median absolute deviation (MAD) values are shown. Two-tailed T-tests showed that T. forsythia was at significantly higher abundance in patients with periodontitis (*Average* ± *s.d.; 4.5* ± *1.4 a.u) than those in health (2.4* ± *0.6 a.u) (P*<*0.05*).

**Table 1.**
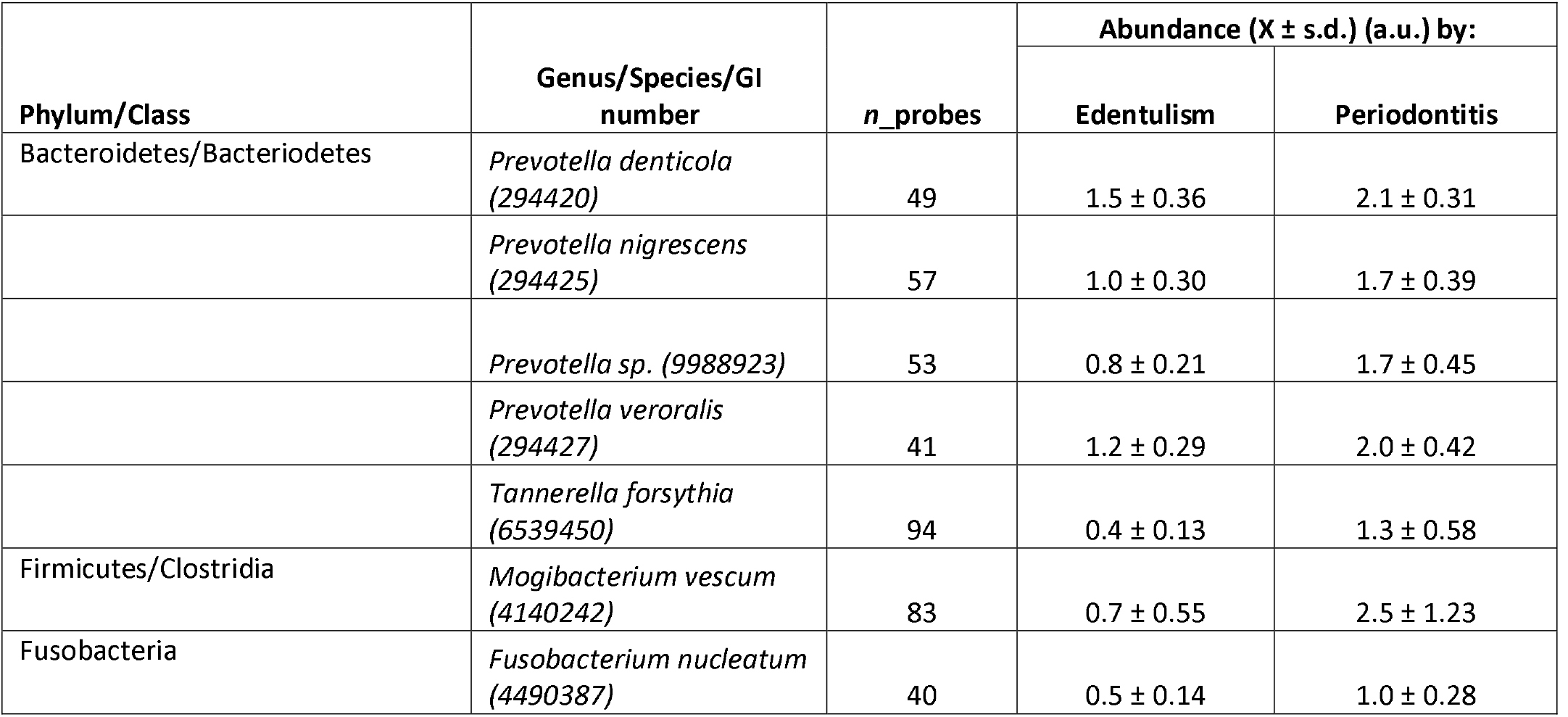
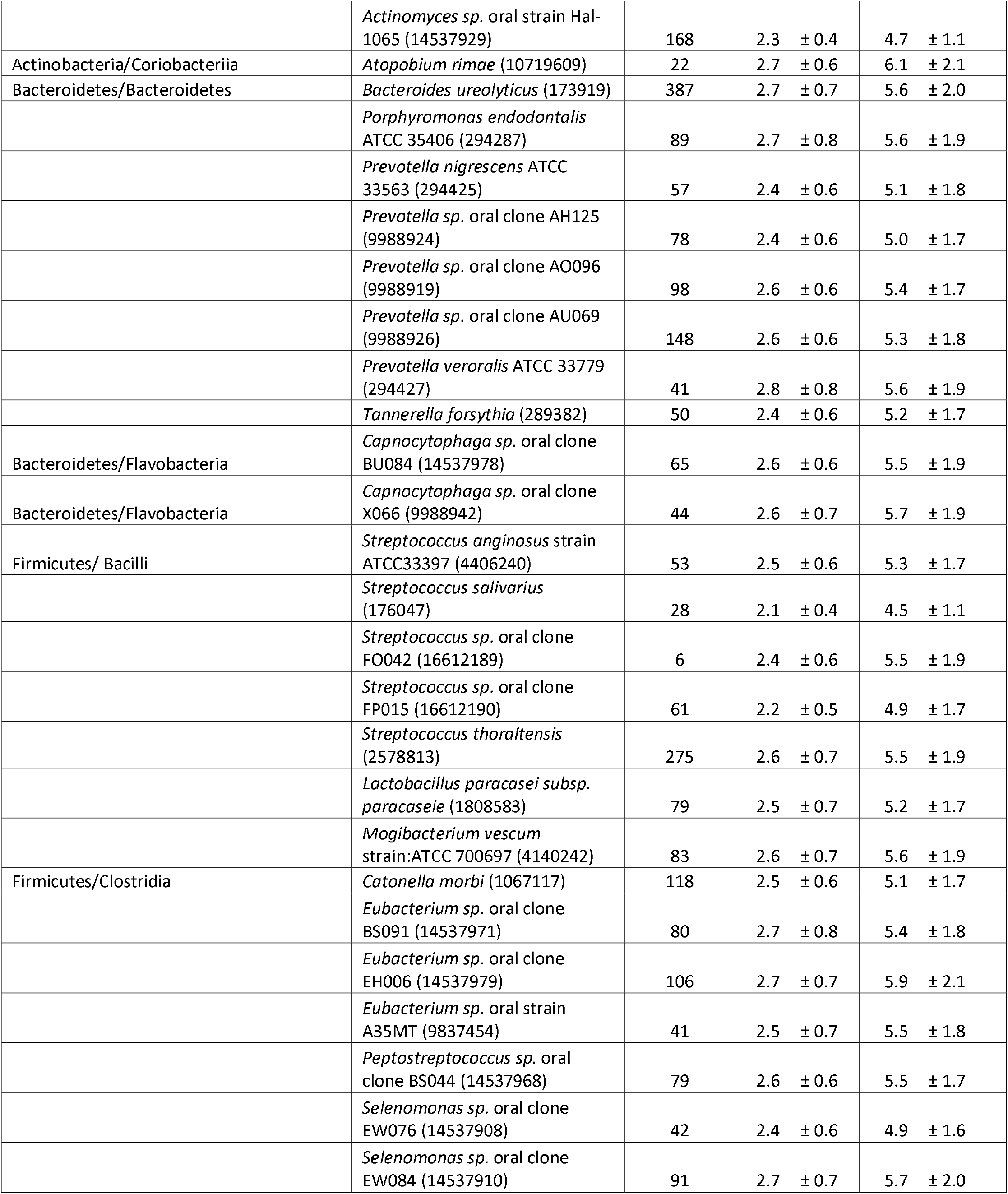
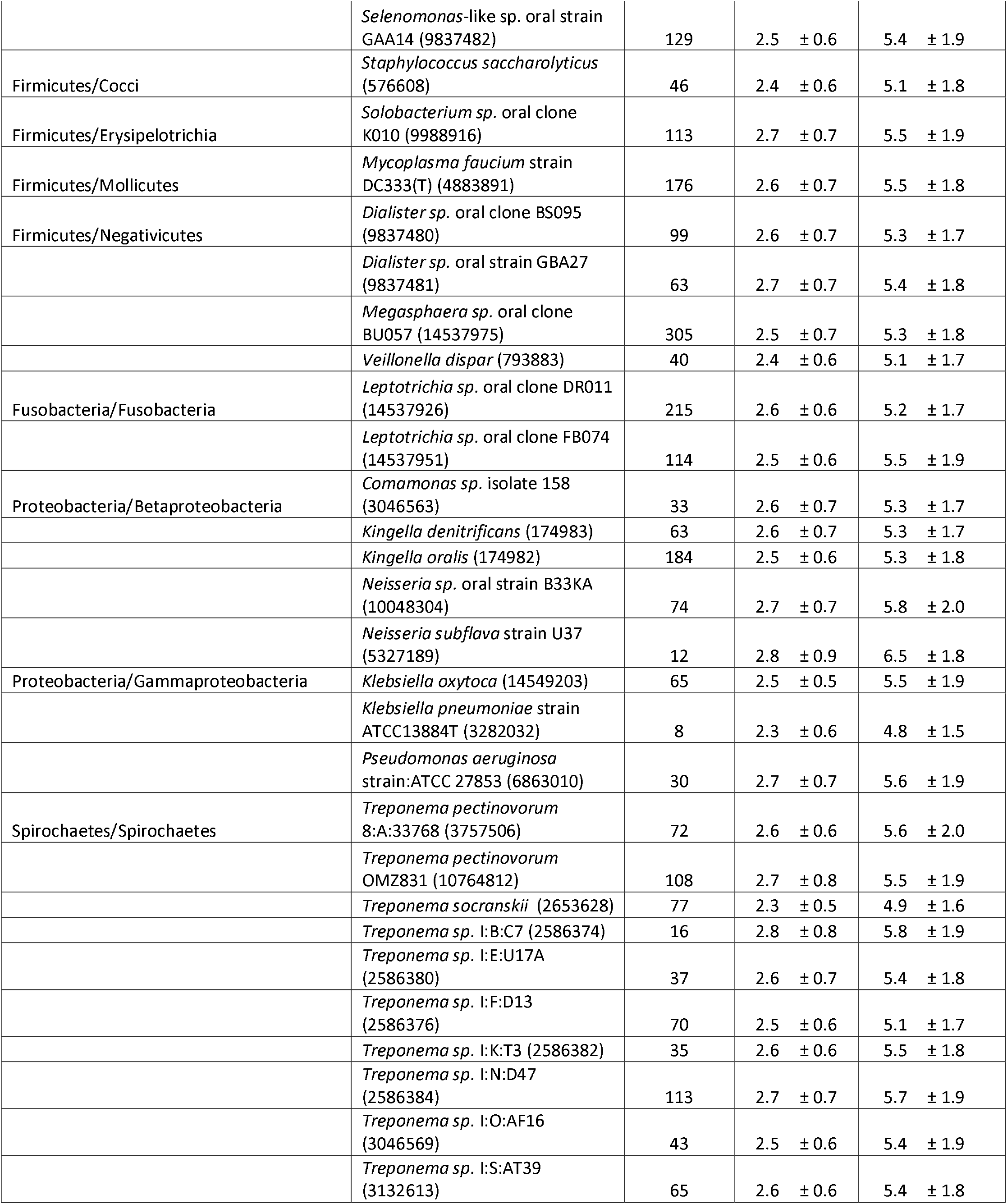
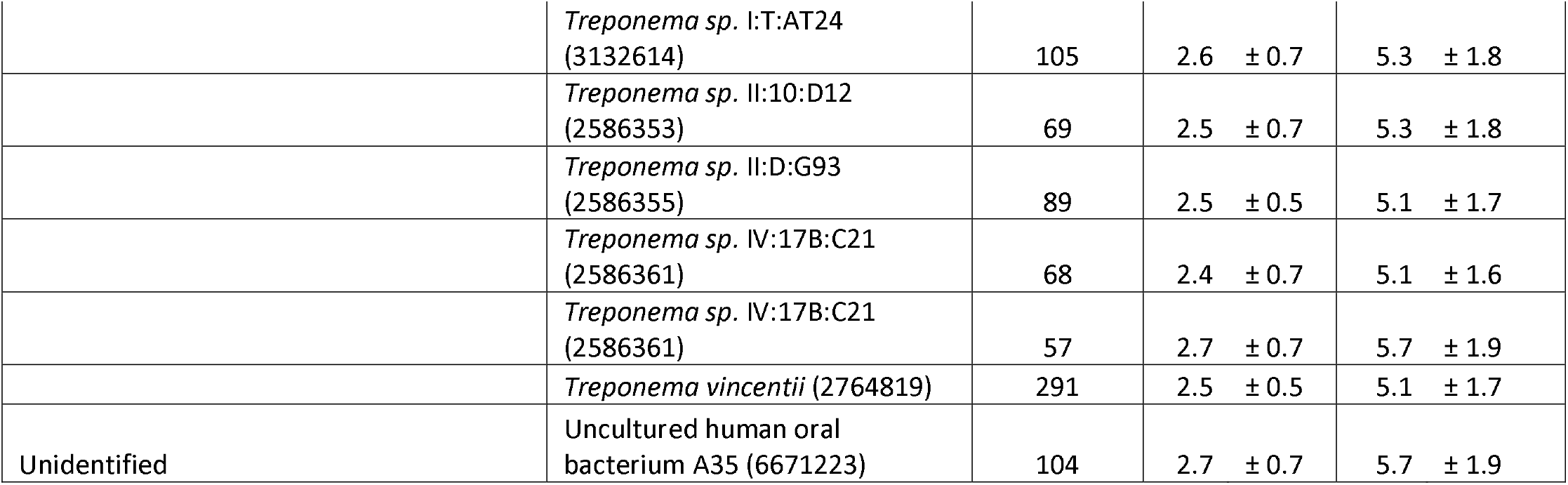
Bacterial species having significantly different abundances for patients with health versus those with edentulism based on individual probes. n_probes, number of probes used to determine the species abundance; Two-sided T-tests were based on four patients with health and four patients with edentulism and alpha = 0.05.

The histograms and whisker plots of the *Treponema denticola* 16S rRNA gene show the distribution of abundances in the 16 patients using 94 unique probes (Figure 3). The average abundances of patients with health, edentulism, periodontitis, and caries were: 2.5 ± 0.6 a.u., 5.3 ± 2.0 a.u., 4.4 ± 1.6 a.u., and 3.8 ± 3.3 a.u., respectively. A two-tailed t-test revealed a difference in average abundances for the 4 patients with health and 4 patients with edentulous but only approached significance (P<=0.06) (Table 2). None of the other paired conditions (i.e., caries versus edentulism, caries versus periodontitis, caries versus health, edentulism versus periodontitis, or health versus periodontitis) for this gene yielded differences from one another.

**Figure 3.**
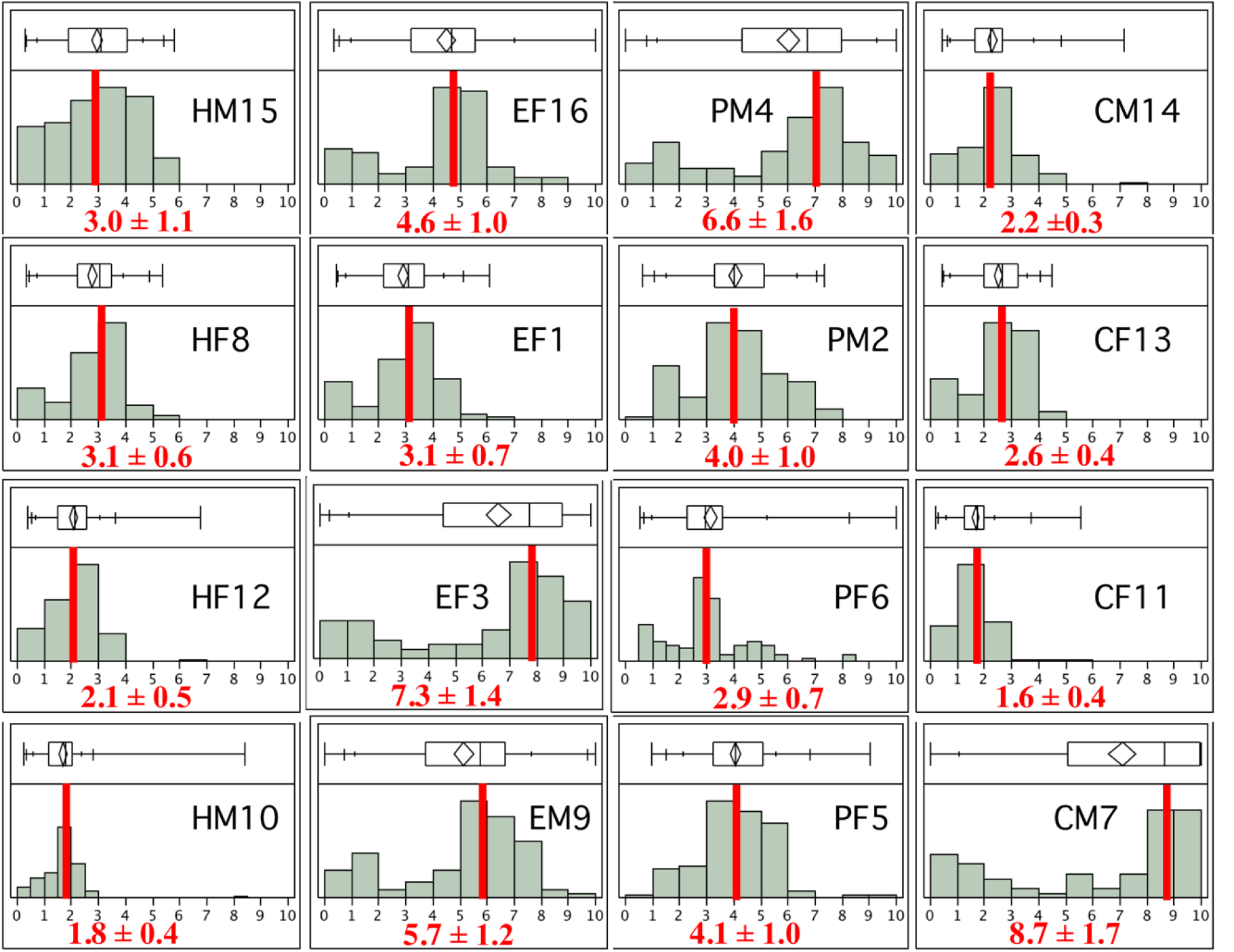
Distribution of 16S rRNA gene abundances for Treponema denticola ATCC 35405 (GI 4580728) in 16 patients based on 94 calibrated probes. Red bar indicates median value. Corresponding MAD values are shown. There is a tendency for T. denticola to be more abundant in patients with periodontitis (*Average* ± *s.d.; 4.4* ± *1.6 a.u) than those with health (2.5* ± *0.6 a.u) (P*<*0.09*).

**Table 2.** Bacterial species having significantly different abundances for patients with health versus those with periodontitis based on calibrated individual probes. n_probes, number of probes used to determine the species abundance; Two-sided T-tests were based on four patients with health and four patients with periodontitis and alpha=0.05.

| Phylum/Class | Genus/Species/GI number | *n_probes* | Abundance ($\bar{X} \pm$ s.d.) (a.u.) by: | |
| --- | --- | --- | --- | --- |
|  |  |  | Health | Periodontitis |
| Actinobacteria/Actinobacteria | *Actinomyces odontolyticus* (853707) | 36 | 2.2 ± 0.5 | 4.1 ± 1.1 |
| | *Propionibacterium avidum* DSM 4901 (2644976) | 53 | 2.7 ± 0.7 | 4.7 ± 1.3 |
| Bacteroidetes/Bacteroidetes | *Prevotella nigrescens* ATCC 33563 (294425) | 57 | 2.4 ± 0.6 | 4.4 ± 1.3 |
| | *Tannerella forsythia* (289382) | 50 | 2.4 ± 0.6 | 4.5 ± 1.4 |
| Bacteroidetes/Flavobacteria | *Capnocytophaga sp.* oral clone X089 (9988944) | 37 | 2.6 ± 0.6 | 4.9 ± 1.4 |
| Firmicutes/Bacilli | *Streptococcus gordonii* (2183315) | 14 | 2.7 ± 0.9 | 4.7 ± 0.9 |
|  | *Streptococcus sp.* oral clone BW009 (9988906) | 11 | 2.7 ± 0.6 | 5.1 ± 1.3 |
|  | *Streptococcus sp.* oral strain T1-E5 (14537933) | 19 | 2.6 ± 0.6 | 4.3 ± 1.1 |
|  | *Streptococcus sp.* oral strain T4-E3 (14537934) | 46 | 2.3 ± 0.5 | 4.3 ± 1.3 |
| Firmicutes/Clostridia | *Selenomonas sp.* oral clone AA024 (9837490) | 96 | 2.6 ± 0.7 | 4.7 ± 1.3 |
| Firmicutes/Cocci | *Staphylococcus capitis* (576605) | 18 | 2.5 ± 0.5 | 4.3 ± 1.1 |
| Fusobacteria | *Fusobacterium periodonticum* | 39 | 2.6 ± 0.7 | 4.7 ± 1.3 |
| Proteobacteria/Betaproteobacteria | *Neisseria flava* strain U40 (5327199) | 1 | 2.7 ± 0.8 | 4.8 ± 0.8 |
|  | *Neisseria sp.* oral clone AP015 (10048301) | 15 | 2.7 ± 0.7 | 5.2 ± 1.4 |
|  | *Neisseria sp.* oral strain B33KA (10048304) | 74 | 2.7 ± 0.7 | 5.1 ± 1.6 |
|  | *Neisseria subflava* (5327189) | 12 | 2.8 ± 0.9 | 5.0 ± 1.3 |
| Proteobacteria/Deltaproteobacteria | *Desulfobulbus sp.* oral clone CH031 (10048312) | 113 | 2.6 ± 0.6 | 4.7 ± 1.4 |
| Proteobacteria/Gammaproteobacteria | *Klebsiella pneumoniae* strain ATCC13884T (3282032) | 8 | 2.3 ± 0.6 | 4.3 ± 0.9 |
|  | *Pseudomonas aeruginosa* strain LMG 1242T (1907091) | 27 | 2.9 ± 0.8 | 5.6 ± 1.8 |
| Spirochaetes/Spirochaetes | *Treponema pectinovorum* 8:A:33768 (3757506) | 72 | 2.6 ± 0.6 | 4.9 ± 1.5 |
|  | *Treponema pectinovorum* OMZ831 (10764812) | 108 | 2.7 ± 0.8 | 4.9 ± 1.5 |
|  | *Treponema socranskii* (2653630) | 35 | 2.4 ± 0.5 | 4.3 ± 1.1 |
|  | *Treponema sp.* II:C:T1 (2586354) | 57 | 2.7 ± 0.7 | 5.1 ± 1.4 |
| Unidentified | Human oral bacterium C20 (6671248) | 2 | 2.3 ± 0.5 | 5.5 ± 1.5 |

### Microbial abundance signatures

To identify microbial abundance signatures (i.e., 16S rRNA genes) that are unique to a particular condition, we compared the 576 rRNA sequences by condition (Caries, Edentulism, Periodontitis and Health) using two-tailed T-tests with unequal variance at alpha=0.05. No significant differences were found for the Caries versus Edentulism, Caries versus Health, Caries versus Periodontitis, or Edentulism versus Periodontitis conditions. However, specific microbial abundance signatures were found in Health versus Edentulism and Health versus Periodontitis conditions (Tables 1 and 2), which are described below.

### Health versus Edentulism

Patients with edentulism had higher abundances of 26 genera and 35 bacterial species than patients associated with dental health. The bacterial species included those in the phyla Actinobacteria, Bacterioidetes, Firmicutes, Fusobacterium, Proteobacteria, and Spirochaetes (Table 1). The number of unique probes used to determine gene abundances of these microorganisms ranged from 6 to 387. Similar to the above, highly related strains (e.g., *Actinomyces odontolyticus* GI 853707 and *A. odontolyticus* GI 69977974) yielded similar abundances even though they were based on different probes (i.e., 36 versus 31 unique probes, respectively). This phenomenon was observed in different strains and species of the genera *Actinomyces, Prevotella, Capnocytophaga, Streptococcus, Eubacterium, Selenomonas, Dialister, Leptotrichia, Kingella, Neisseria, Klebsiella*, and *Treponema*.

### Health versus periodontitis

Patients with periodontitis had higher abundances of 14 genera and 21 bacterial species than patients with health. The bacterial genera included those found in the phyla Actinobacteria, Bacterioidetes, Firmicutes, Fusobacterium, Proteobacteria, and Spirochaetes (Table 2). Note that the number of unique probes used to determine gene abundances varied from one to 113. We emphasize this point because additional probes targeting a particular 16S rRNA gene presumably improved the precision. Interestingly, strains of highly related species (e.g., *Streptococcus sp.*; GI 14537933 and GI 14537934) yielded similar abundances even though they were based on different probes (i.e., 19 versus 46 unique probes, respectively). This phenomenon was also observed in different strains and species of the genera *Streptococcus, Neisseria* and *Treponema*. This finding provides support for the precision of Gene Meters in terms of gene abundances.

### Calibrated probe aggregates

The PMT settings affected the number of total genes (*n*=597) that could be calibrated. Specifically, 564, 567, and 572 genes were calibrated for the PMT settings of 500, 600 and 700, respectively. The abundances for these genes in the 16 patient samples were assembled into a dataset by PMT (Table S3). As before, gene abundances that were greater than 10 a.u. were set to 10 a.u. because we did not expect calibration to forecast accurately beyond the dilution series used to create it. The average gene abundance for the 500, 600, and 700 datasets was 2.9 ± 1.8, 3.4 ± 1.9, and 4.0 ± 2.1 arbitrary units (a.u.), respectively. The lowest and highest gene abundances for all datasets was 0.03 and 10.0 a.u., respectively. Therefore, gene abundances could vary by as much as 300-fold and the averaged gene abundances were affected by PMT settings, with higher abundances at higher PMT settings. Based on these results, all subsequent analyses of the aggregated calibrated probes were performed using the 600 PMT setting data.

### Microbial abundance signatures

Microbial abundance signatures (i.e., 16S rRNA genes) were determined by comparing the 567 rRNA sequences by condition (Caries, Edentulism, Periodontitis and Health) using two-tailed T-tests with unequal variance at alpha=0.05. Although significant differences were not found for the Caries versus Health comparison, specific microbial abundance signatures were found for the Caries versus Edentulism, Caries versus Periodontitis, Edentulism versus Health, Edentulism versus Periodontitis, and Health versus Periodontitis comparisons, which are described below.

### Caries versus Edentulism

Patients with edentulism had significantly higher abundances of *Lactobacillus sp.* (2.8 ± 1.21 a.u.) than patients with caries (0.6 ± 0.42 a.u.) (*P*<0.03). Hence, *Lactobacillus sp.* was a putative abundance signature for patients with edentulism.

### Caries versus Periodontitis

Patients with periodontitis had significantly higher abundances of *Tannerella forsythia* (1.3 ± 0.58 a.u.) than patients with caries (0.4 ± 0.28 a.u.) (*P*<0.04). Similarly, patients with periodontitis had significantly higher abundances of *Fusobacterium nucleatum* (1.0 ± 0.28 a.u.) than patients with caries (0.4 ± 0.22 a.u.) (*P*<0.01). The high abundances of these microorganisms indicates a putative signature for periodontitis as these microorganisms were significantly more abundant in patients with periodontitis versus patients with health (see below).

### Edentulism versus Health

Patients with edentulism had significantly higher abundances of 32 genera (and two unidentified species) representing 72 different species/strains than patients associated with health. The microorganisms included the phyla Actinobacteria, Ascomycota, Bacterioidetes, Deinococcus-Thermus, Firmicutes, Fusobacterium, Proteobacteria, Spirochaetes, and Tenericutes (Table 3). The phylum Firmicutes contained the most microorganisms (*n*=29 species) with the following classes: Bacilli, Clostridia, Erysipelotrichia, and Negativicutes. The class Bacilli consisted of *Bacillus clausii*, and several *Lactobacillus* and *Streptococcus* species. The phylum Actinobacteria contained the second most microorganisms (*n*=15 species) and included the classes Actinobacteria and Coriobacterila. Within the class Actinobacteria were the following genera: *Actinomyces, Bifidobacterium, Propionibacterium*, and *Stomatococcus*.

**Table 3.** Microbial species having significantly different abundances for patients with health versus those with edentulism using calibrated aggregated probes. n_probes, number of probes used to determine the species abundance; Two-sided T-tests were based on four patients with health and four patients with edentulism and alpha=0.05.

| Phylum/Class | Genus/Species/GI number | $n\_probes$ | Abundance ($\bar{X} \pm s.d.$) (a.u.) by: | |
| --- | --- | --- | --- | --- |
|  |  |  | Health | Edentulism |
| Actinobacteria/Actinobacteria | *Actinomyces gerencseriae* (1838939) | 193 | $1.9 \pm 0.36$ | $3.1 \pm 0.80$ |
| | *Actinomyces israelii* (1838944) | 314 | $1.6 \pm 0.26$ | $2.7 \pm 0.67$ |
| | *Actinomyces meyeri* (1838945) | 74 | $2.1 \pm 0.41$ | $4.4 \pm 1.42$ |
| | *Actinomyces odontolyticus* (6997974) | 31 | $2.5 \pm 0.48$ | $5.4 \pm 1.73$ |
| | *Actinomyces odontolyticus* (853707) | 36 | $1.6 \pm 0.32$ | $2.7 \pm 0.54$ |
| | *Actinomyces sp.* (14537925) | 310 | $1.7 \pm 0.31$ | $3.0 \pm 0.86$ |
| | *Actinomyces sp.* (14537930) | 56 | $2.4 \pm 0.52$ | $5.0 \pm 1.64$ |
| | *Actinomyces sp.* (14537912) | 82 | $2.0 \pm 0.42$ | $4.0 \pm 1.26$ |
| | *Actinomyces sp.* (14537929) | 168 | $1.8 \pm 0.30$ | $3.5 \pm 0.32$ |
| | *Actinomyces sp.* (2073386) | 131 | $1.6 \pm 0.21$ | $2.6 \pm 0.62$ |
| | *Actinomyces viscosus* (1838951) | 98 | $2.2 \pm 0.39$ | $3.7 \pm 0.93$ |
|  | *Bifidobacterium sp. (9837452)* | 205 | 2.0 ± 0.33 | 3.4 ± 0.89 |
|  | *Propionibacterium avidum (2644976)* | 53 | 2.5 ± 0.54 | 5.4 ± 1.82 |
|  | *Stomatococcus mucilaginosus (2764819)* | 291 | 1.8 ± 0.17 | 2.8 ± 0.34 |
| Actinobacteria/Coriobacteriia | *Atopobium rimae (10719609)* | 22 | 2.7 ± 0.62 | 7.6 ± 2.13 |
| Ascomycota/Saccharomycetes | *Candida albicans (2507)* | 1442 | 2.2 ± 0.46 | 4.7 ± 1.56 |
| Bacteroidetes/Bacteroidetes | *Porphyromonas-like sp. (9988935)* | 631 | 1.9 ± 0.31 | 3.1 ± 0.82 |
|  | *Prevotella sp. (9988921)* | 87 | 2.6 ± 0.56 | 5.6 ± 1.94 |
|  | *Prevotella sp. (14537919)* | 88 | 2.6 ± 0.60 | 5.7 ± 2.02 |
|  | *Prevotella sp. (9988918)* | 127 | 1.9 ± 0.33 | 3.2 ± 0.85 |
|  | *Prevotella sp. (9988919)* | 98 | 2.2 ± 0.37 | 3.7 ± 0.97 |
| Bacteroidetes/Flavobacteria | *Capnocytophaga sp. (9988943)* | 101 | 2.7 ± 0.62 | 5.9 ± 2.08 |
| Bacteroidetes/Sphingobacteria | *Pedobacter sp. (14537940)* | 418 | 2.2 ± 0.44 | 4.4 ± 1.39 |
| Deinococcus-Thermus/Deinococci | *Deinococcus sp. (14537950)* | 790 | 2.1 ± 0.43 | 4.2 ± 1.39 |
| Firmicutes/Bacilli | *Bacillus clausii (9715774)* | 27 | 2.2 ± 0.43 | 4.7 ± 1.57 |
|  | *Lactobacillus gasseri (175028)* | 309 | 2.1 ± 0.45 | 4.5 ± 1.54 |
|  | *Lactobacillus paracasei (1808583)* | 79 | 1.7 ± 0.32 | 2.7 ± 0.71 |
|  | *Lactobacillus sp. (9988912)* | 73 | 0.5 ± 0.13 | 2.8 ± 1.21 |
|  | *Paenibacillus sp. (15004607)* | 149 | 2.0 ± 0.35 | 3.6 ± 1.04 |
|  | *Streptococcus gordonii (2183315)* | 14 | 1.8 ± 0.33 | 2.8 ± 0.65 |
|  | *Streptococcus mitis (9988909)* | 6 | 2.7 ± 0.80 | 5.7 ± 1.95 |
|  | *Streptococcus pneumoniae (2183314)* | 29 | 1.9 ± 0.40 | 4.1 ± 1.33 |
|  | *Streptococcus salivarius (176047)* | 28 | 1.9 ± 0.40 | 4.1 ± 1.10 |
|  | *Streptococcus sp. (9988907)* | 91 | 2.5 ± 0.60 | 5.4 ± 1.87 |
|  | *Streptococcus sp. (16612185)* | 83 | 2.1 ± 0.45 | 4.3 ± 1.38 |
|  | *Streptococcus thoraltensis (2578813)* | 275 | 2.3 ± 0.49 | 4.9 ± 1.60 |
| Firmicutes/Clostridia | *Butyrivibrio sp. (9837461)* | 569 | 1.8 ± 0.35 | 3.5 ± 1.10 |
|  | *Eubacterium limosum (174517)* | 437 | 2.1 ± 0.58 | 4.5 ± 1.58 |
|  | *Eubacterium minutum* (4001750) | 60 | 1.2 ± 0.21 | 2.0 ± 0.51 |
|  | *Eubacterium saphenum*<br>(1513214) | 447 | 1.7 ± 0.30 | 3.0 ± 0.77 |
|  | *Eubacterium* sp. (14537976) | 196 | 2.1 ± 0.37 | 3.6 ± 0.96 |
|  | *Eubacterium* sp. (14537971) | 80 | 1.5 ± 0.43 | 2.9 ± 0.87 |
|  | *Eubacterium* sp. (14537961) | 219 | 1.9 ± 0.32 | 3.4 ± 0.88 |
|  | *Peptostreptococcus anaerobius*<br>(175621) | 132 | 1.7 ± 0.28 | 2.9 ± 0.78 |
|  | *Peptostreptococcus*<br>*asaccharolyticus* (454180) | 439 | 1.9 ± 0.42 | 3.5 ± 0.98 |
|  | *Eubacterium* sp. (3861469) | 515 | 2.0 ± 0.35 | 3.3 ± 0.87 |
| Firmicutes/Erysipelotrichia | *Erysipelothrix tonsillarum*<br>(7678893) | 478 | 1.8 ± 0.34 | 3.4 ± 1.03 |
|  | *Solobacterium* sp. (9988916) | 113 | 1.9 ± 0.39 | 3.5 ± 1.08 |
| Firmicutes/Negativicutes | *Selenomonas flueggei* (9837496) | 95 | 1.7 ± 0.35 | 3.3 ± 1.05 |
|  | *Selenomonas flueggei* (9837497) | 50 | 2.2 ± 0.53 | 4.8 ± 1.50 |
|  | *Uncultured Selenomonas*<br>(14161358) | 84 | 2.7 ± 0.70 | 5.8 ± 2.00 |
|  | *Uncultured Veillonella*<br>(14161350) | 47 | 1.7 ± 0.44 | 2.9 ± 0.80 |
|  | *Veillonella dispar* (793883) | 40 | 1.8 ± 0.34 | 3.4 ± 0.81 |
| Fusobacteria/Fusobacteria | *Leptotrichia* sp. (16612191) | 400 | 2.4 ± 0.57 | 5.0 ± 1.69 |
|  | *Leptotrichia* sp. (14537926) | 215 | 2.4 ± 0.50 | 4.7 ± 1.48 |
| Proteobacteria/Betaproteobacteria | *Burkholderia* sp. (9988903) | 264 | 2.0 ± 0.35 | 3.3 ± 0.85 |
|  | *Kingella denitrificans* (174983) | 63 | 2.2 ± 0.36 | 3.6 ± 0.91 |
|  | *Neisseria* sp. (10048304) | 74 | 2.7 ± 0.57 | 5.9 ± 2.05 |
| Proteobacteria/Epsilonproteobacteria | *Campylobacter* sp. (10048314) | 239 | 1.7 ± 0.33 | 3.0 ± 0.84 |
| Proteobacteria/Gammaproteobacteria | *Ewingella americana* (903619) | 272 | 2.0 ± 0.39 | 4.0 ± 1.30 |
|  | *Klebsiella pneumoniae* (3282032) | 8 | 2.2 ± 0.53 | 4.8 ± 1.61 |
| Spirochaetes/Spirochaetes | *Treponema denticola* (4580728) | 94 | 1.0 ± 0.16 | 1.7 ± 0.41 |
|  | *Treponema pectinovorum*<br>(10764812) | 108 | 2.6 ± 0.64 | 5.6 ± 1.95 |
|  | *Treponema socranskii* (2653628) | 77 | 2.3 ± 0.56 | 4.7 ± 1.52 |
|  | *Treponema sp.* (3132614) | 105 | 2.7 ± 0.51 | 4.5 ± 1.50 |
|  | *Treponema sp.* (2586354) | 57 | 2.6 ± 0.65 | 5.7 ± 1.98 |
|  | *Treponema sp.* (14537906) | 128 | 0.9 ± 0.16 | 1.5 ± 0.43 |
|  | *Treponema sp.* (6049684) | 85 | 2.7 ± 0.71 | 6.0 ± 2.04 |
|  | *Treponema sp.* (2586355) | 89 | 1.4 ± 0.21 | 2.3 ± 0.56 |
| Tenericutes/Mollicutes | *Mycoplasma buccale* (4883887) | 191 | 2.6 ± 0.61 | 5.8 ± 1.95 |
| Unknown | Human oral (6671248) | 2 | 2.7 ± 0.60 | 4.9 ± 1.29 |
|  | Uncultured human (6671223) | 104 | 2.6 ± 0.64 | 5.9 ± 1.91 |

### Edentulism and Periodontitis

Patients with periodontitis had significantly higher abundances of seven microbial species/strains than patients with edentulism (Table 4). The phyla for these bacteria included Bacteroidetes, Firmicutes, and Fusobacteria. Notable genera included: *Prevotella*, *Tannerella*, and *Fusobacterium*, which are thought to play roles in periodontitis.

**Table 4.**
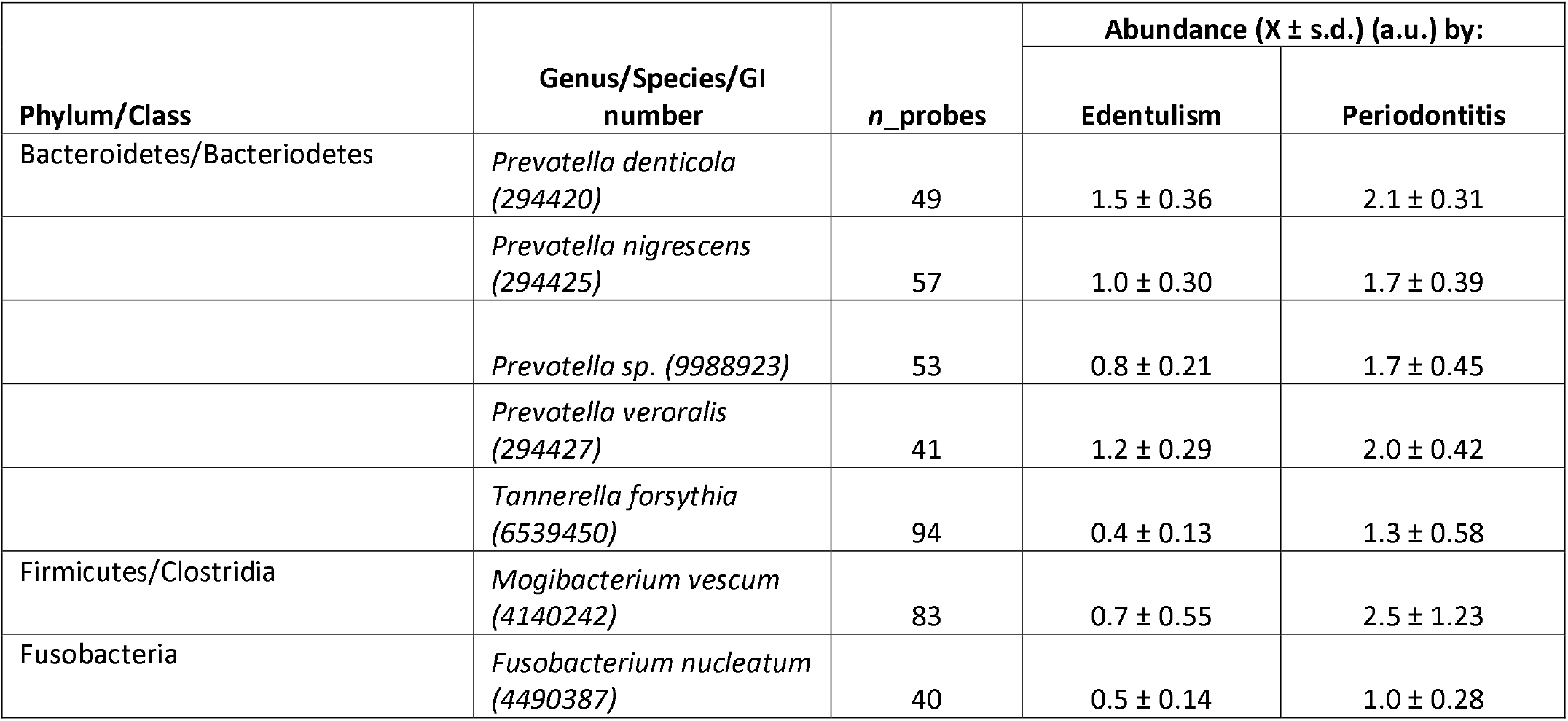
Microbial species having significantly different abundances for patients with edentulism versus those with periodontitis based on aggregated probes. n_probes, number of probes used to determine the species abundance; Two-sided T-tests were based on four patients with edentulism and four patients with periodontitis and alpha=0.05.

### Health and Periodontitis

Patients with periodontitis had higher abundances of 30 genera (five not taxonomically identified) and 62 species than patients with health (Table 5). These bacteria were found in the phyla Actinobacteria, Bacteroidetes, Chloroflexi, Deferribacteres, Firmicutes, Fusobacteria, Proteobacteria, and Spirochaetes. Note that there was a higher abundance of members of the “red complex” (specifically, *Porphyromonas gingivalis, Tannerella forsythia, Treponema denticola*) in patients with periodontitis than patients with health.

**Table 5.**
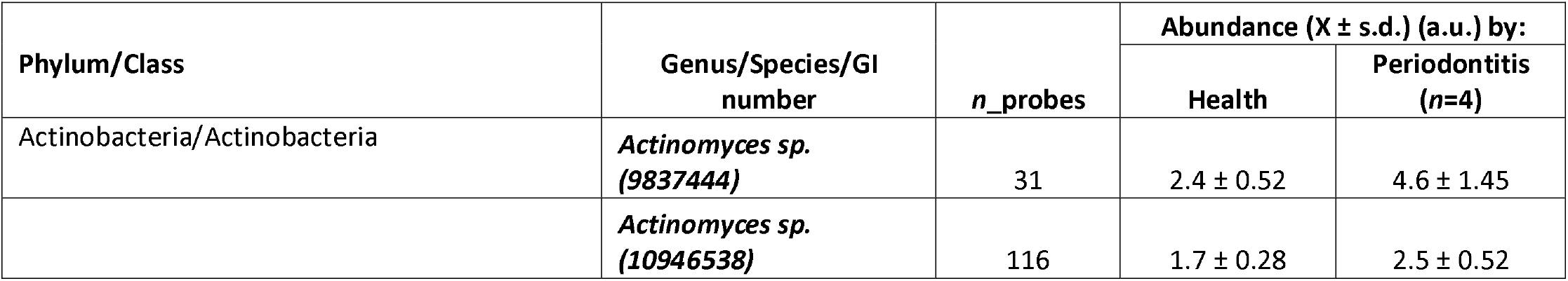
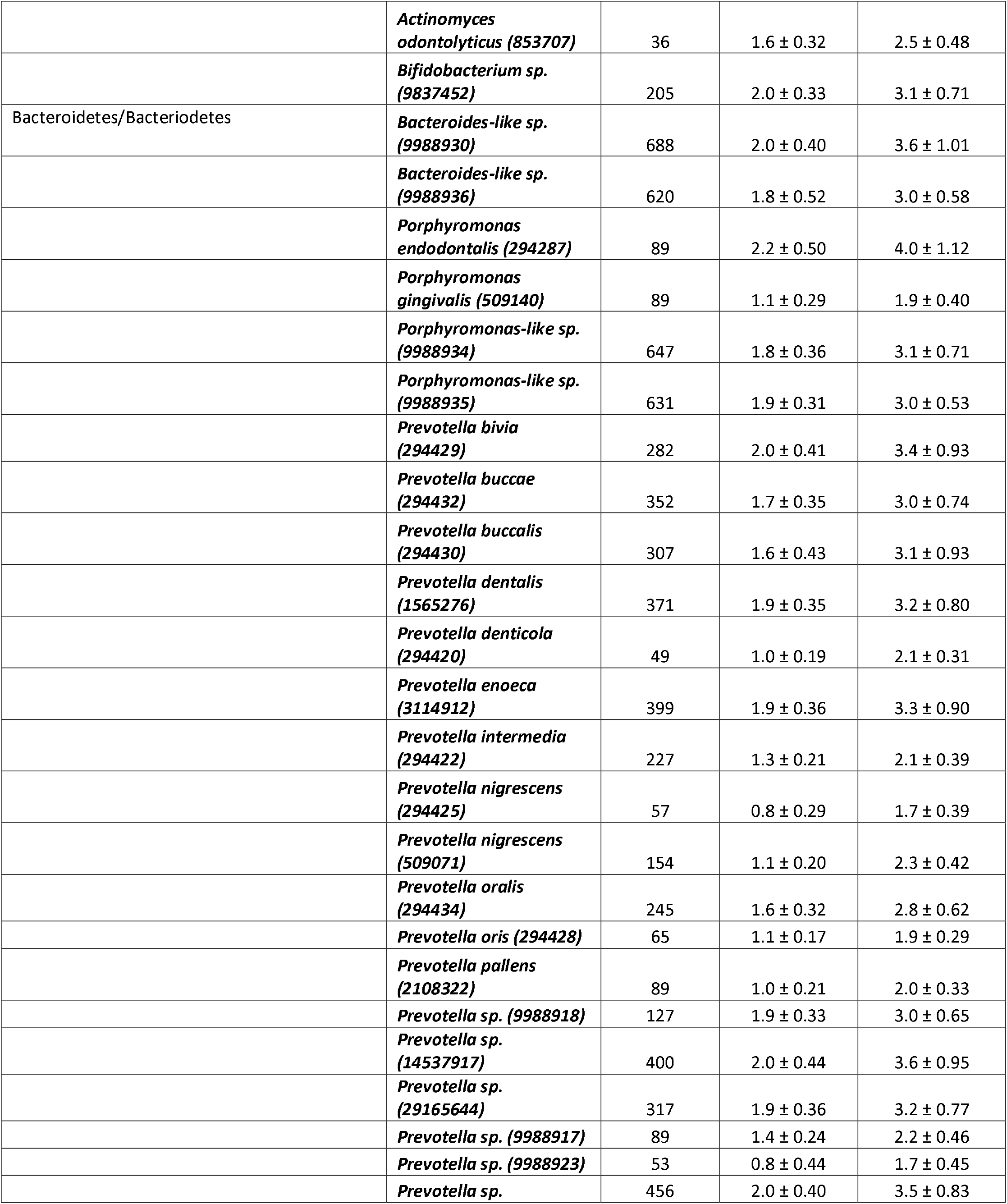
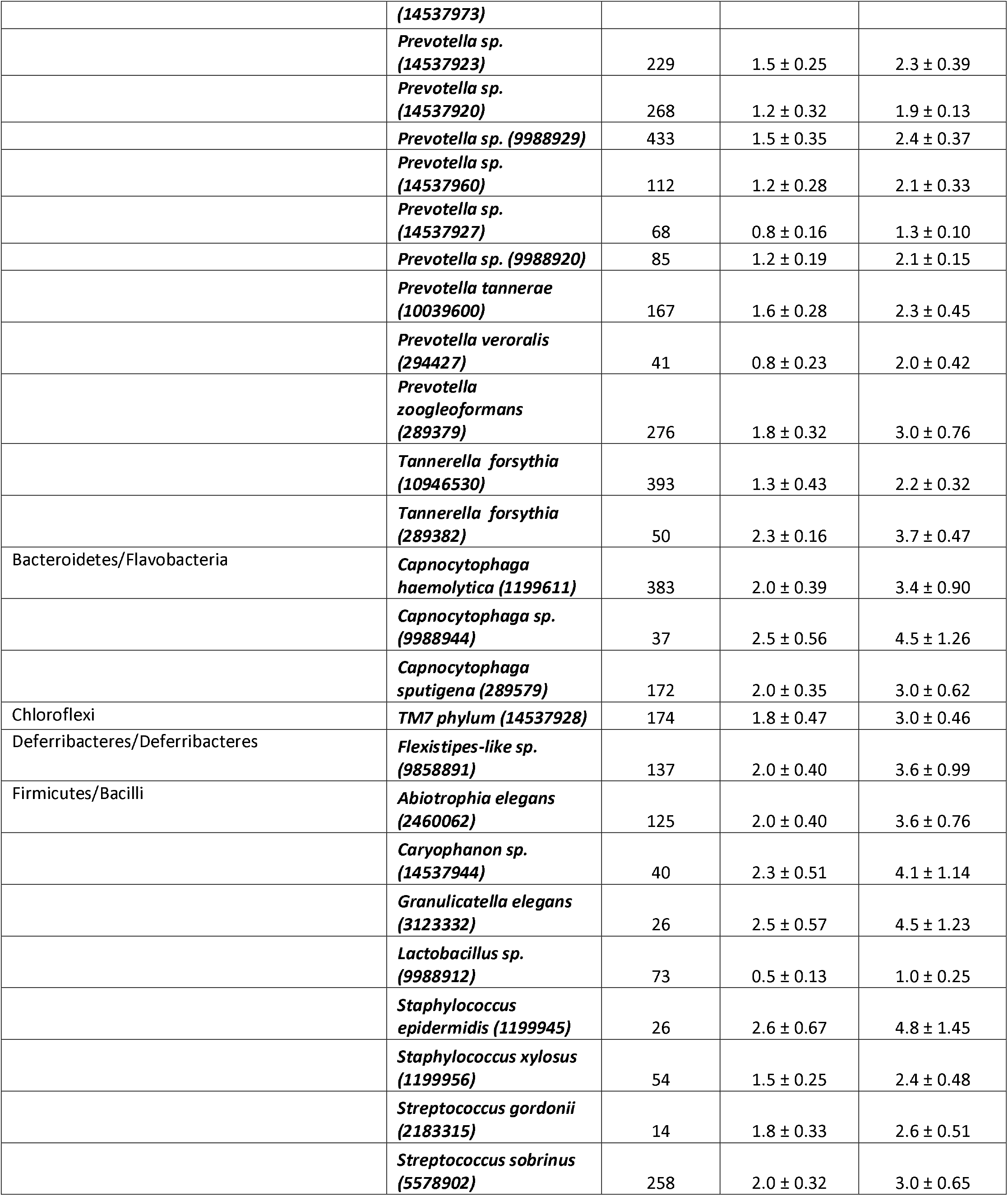
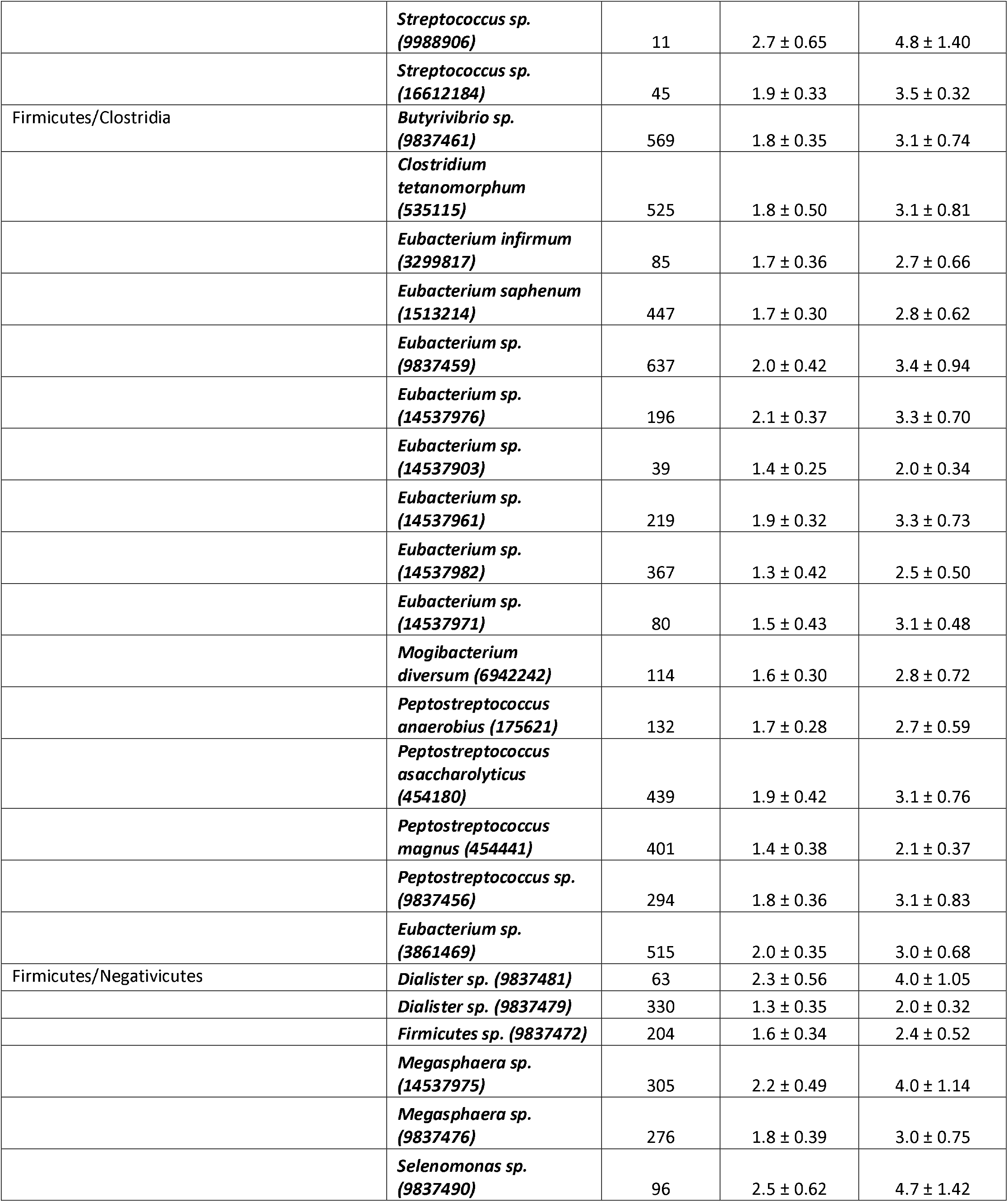
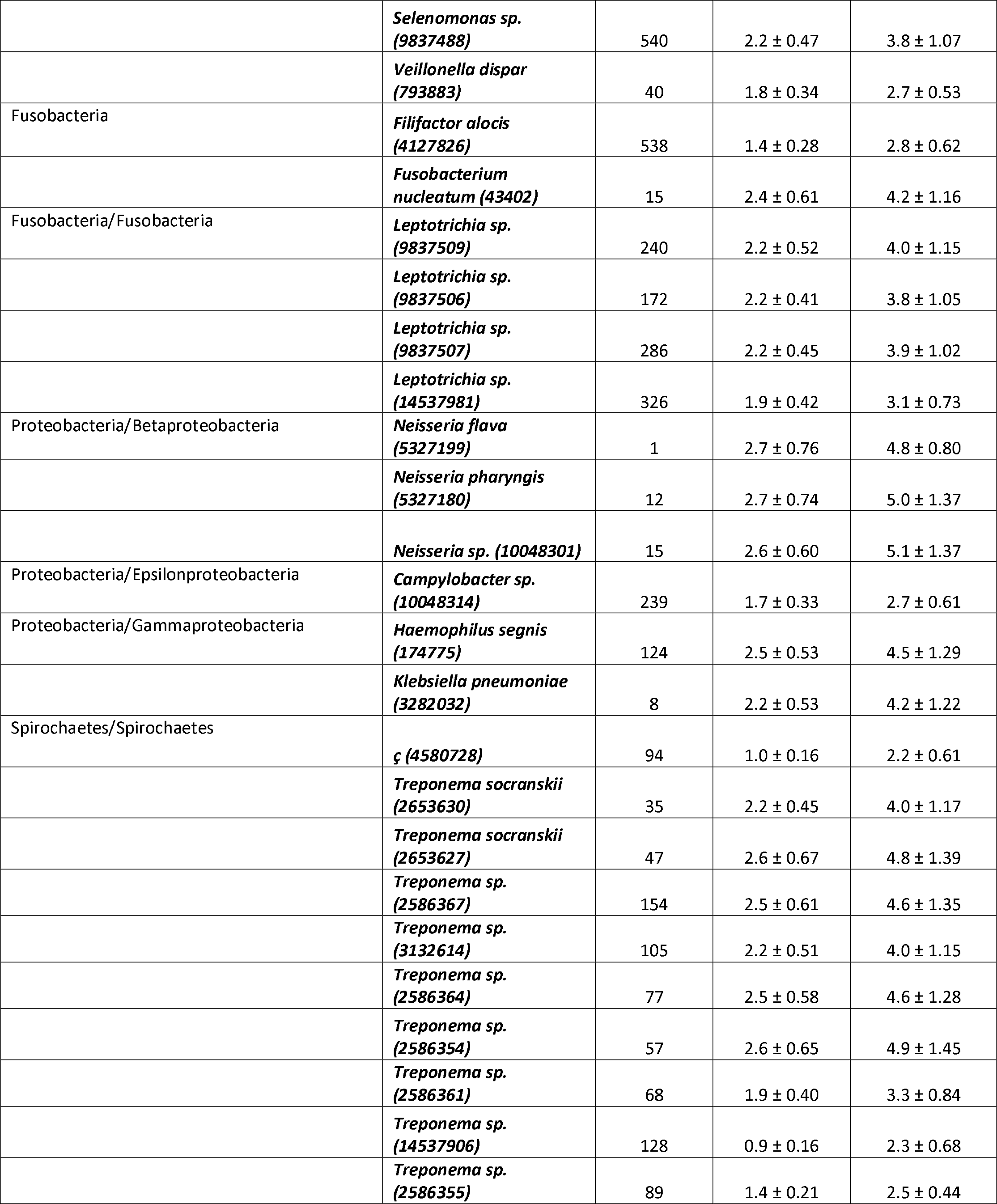
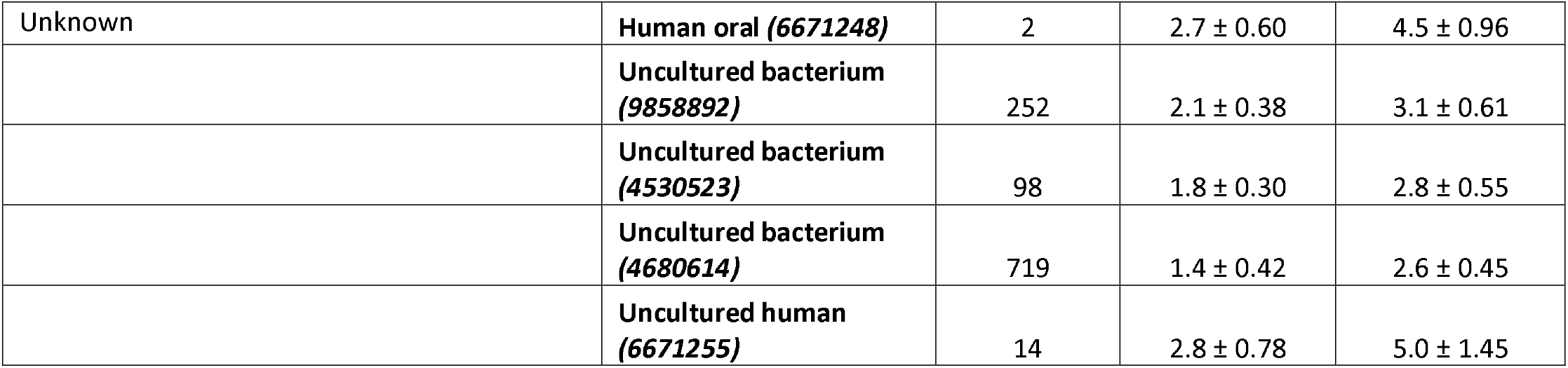
Microbial species having significantly different abundances for patients with health versus those with periodontitis based on aggregated probes. n_probes, number of probes used to determine the species abundance; Two-sided T-tests were based on four patients with health and four patients with periodontitis and alpha=0.05.

### Comparison of calibration methods

Correlation analysis was used to compare calibration methods. While in theory the abundances should be similar by calibration method and the correlations close to one, the abundances determined by aggregated probes were anticipated to be more robust than individual probes when the gene target abundances approached zero. The reason for this is that the sum of all signal intensities obtained from multiple probes targeting the same gene will always be greater than the signal intensities of individual probes. At low gene target abundances, the signal intensities of individually calibrated probes approach the resolution of the scanner, which means the potential for noise in the signal would be minimized in the aggregated probe results.

A plot of the relationship between the correlations of the calibration methods against the average target abundances is shown in Figure 4. While only 65 out of the possible 567 (∼11.5%) microorganisms (GIs) had correlations of less than 0.95, most microorganisms (*n*=502) (i.e., GIs) had similar abundances (∼88.5% had correlation >=0.95). Figure 4 shows that when the average abundance was less than 2 a.u., the correlations were less than 0.95 (Figure 4). The significance of this finding is that it provides support that the aggregate probe calibration is more precise than the individual probe calibration.

**Figure 4.**
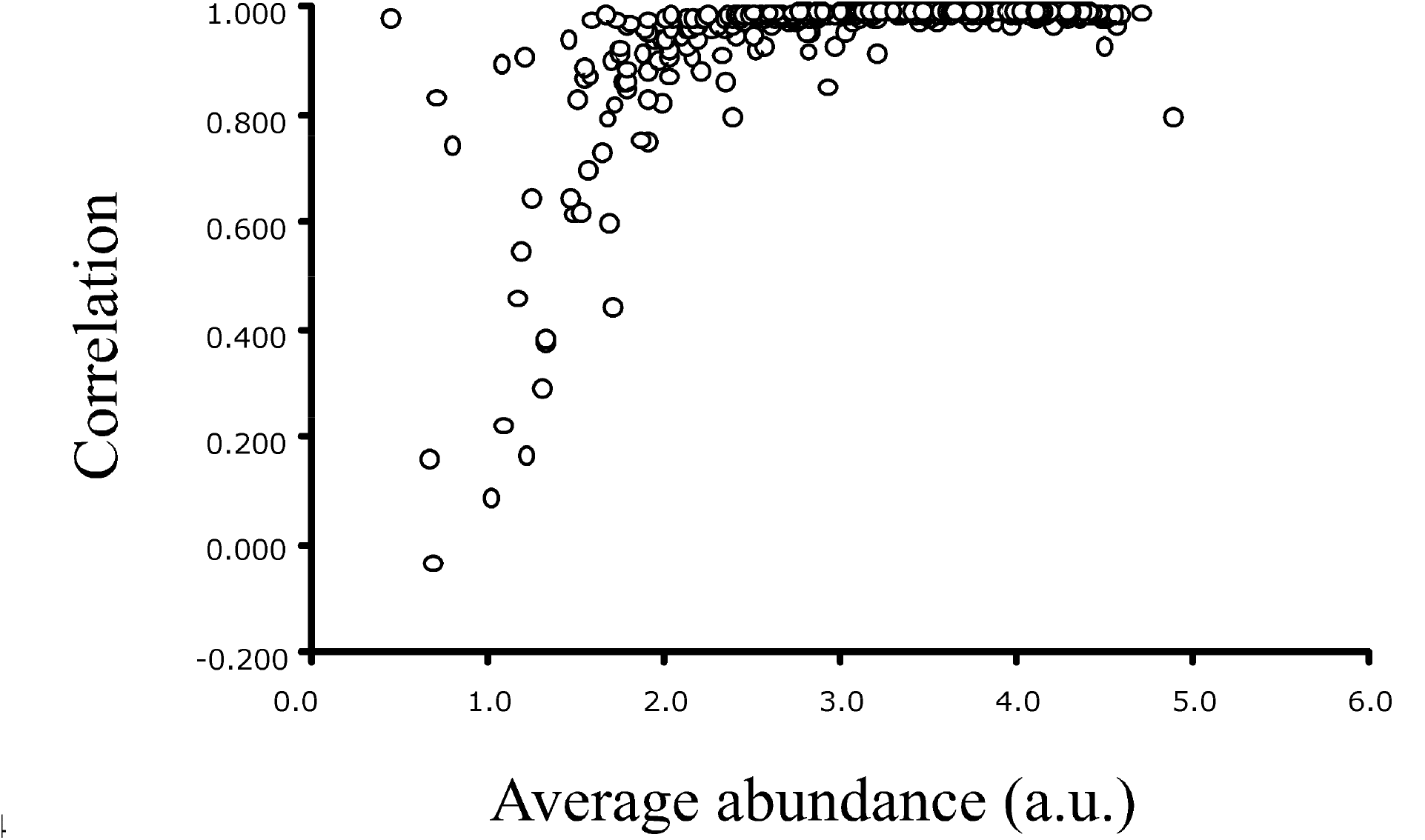
Correlation of abundances of GIs for two calibration approaches (individual probe calibration and aggregate probes targeting same gene) versus average abundance of the GIs determined using the aggregate calibration approach. Results show the average abundances for GIs are low (< 2.0 a.u.). The correlations are also low, presumably because the signal intensities of the individual probes approach the resolution of the scanner while the sum of the signal intensities of aggregated probes do not.

### Gene Meter versus DNA sequencing results

In theory, one would expect the same microorganisms to be identified by both approaches (Gene Meters and DNA sequencing) since the source of DNA was the same PCR amplification products from the same 16 patient samples.

For the Gene Meter approach, we detected 576 microbial targets (564 non-redundant targets) (Figure 5). Thirty-five of the 564 non-redundant targets could not be taxonomically resolved: 13 were classified as “unculturable” and 22 could not be identified to the genus level. Subtracting these 35 ambiguous targets from the 564 non-redundant gene targets left 107 microbial genera.

**Figure 5.**
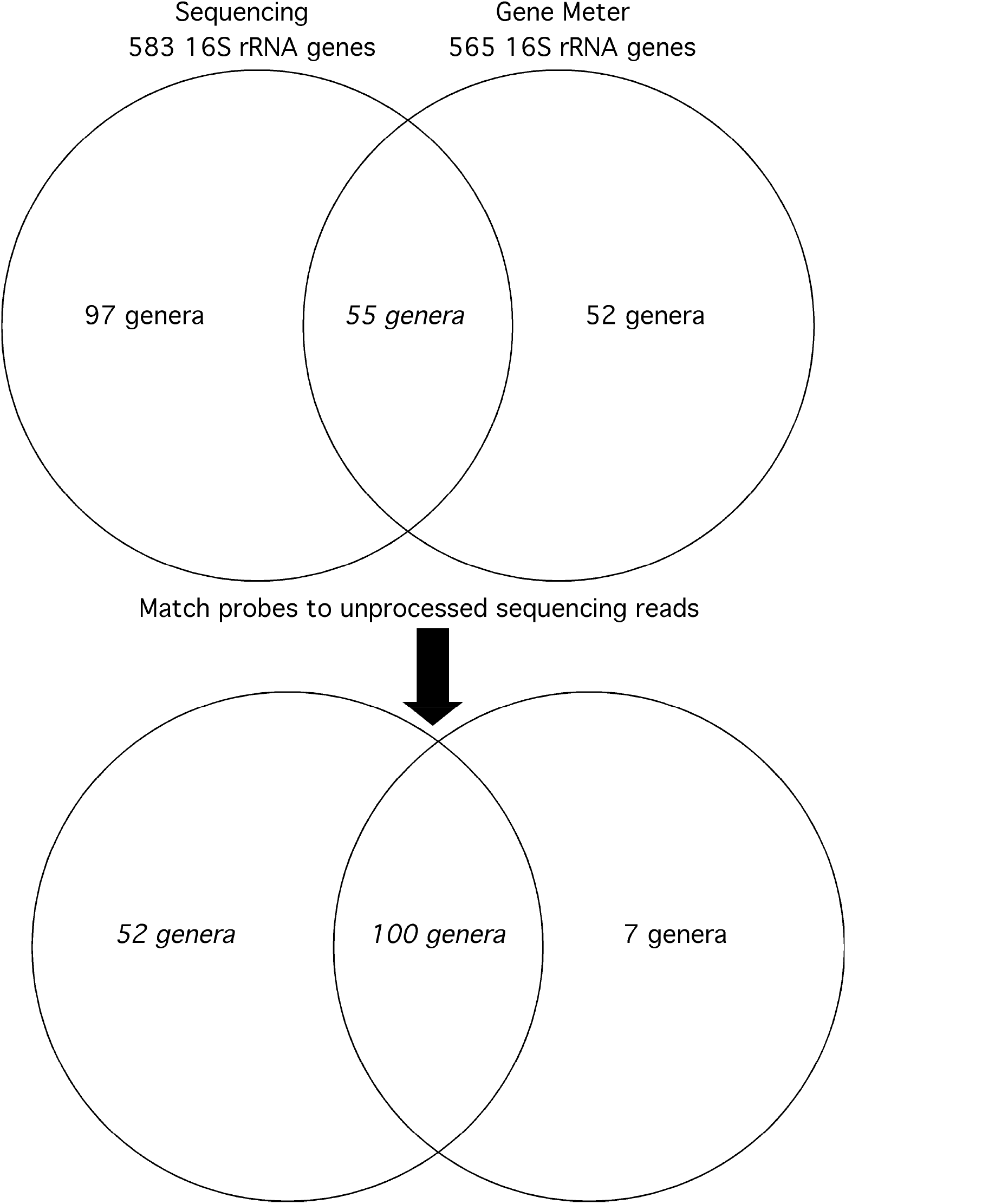
Shared genera by DNA sequencing and Gene Meter for 16 patients. Top shows taxonomic classification based on processed DNA sequencing reads. Bottom shows taxonomic classification based on probe matches of unprocessed DNA sequencing reads and processed DNA sequencing reads.

For the DNA sequencing approach, we detected 654 targets (583 non-redundant targets) (Figure 5). Nine of the 583 non-redundant targets yielded ambiguous taxonomy (e.g., “anaerobic bacterium”, “rumen bacterium”) and 17 were plants (e.g., *Nicotiana sylvestris*, *N. tabacum, N. tomentosiformism,* “Onion yellows”, *Oryza nivara, O. sativa,* and *Phalaenopsis aphrodite*, *Citrus sinensis, Cucumis sativus, Dioscorea elephantipes, Draba nemorosa, Drimys granadensis, Heliathus annuus, Calycanthus floridus, Agrostis stolonifera, Amborella trichopoda, Calycanthus floridus, Gossypium barbadense,* and *Gossypium barbadense*). Subtracting these 26 ambiguous targets from the 583 non-redundant gene targets left 152 microbial genera.

The union of the two approaches yielded 55 common microbial genera: *Abiotrophia, Actinobacillus, Actinomyces, Aggregatibacter, Atopobium, Bacillus, Bacteroides, Bergeyella, Bifidobacterium, Bradyrhizobium, Burkholderia, Butyrivibrio, Campylobacter, Capnocytophaga, Clostridium, Corynebacterium, Cryptobacterium, Desulfovibrio, Dialister, Eikenella, Enterococcus, Erysipelothrix, Eubacterium, Fusobacterium, Gemella, Granulicatella, Haemophilus, Kingella, Lactobacillus, Leptotrichia, Megasphaera, Microbacterium, Micrococcus, Mitsuokella, Moraxella, Mycobacterium, Mycoplasma, Neisseria, Paenibacillus, Peptococcus, Peptostreptococcus, Porphyromonas, Prevotella, Propionibacterium, Pseudomonas, Rothia, Selenomonas, Slackia, Staphylococcus, Streptococcus, Tannerella, Treponema, Variovorax, Veillonella,* and *Xanthomonas.*

Fifty-two of the 107 genera (∼49%) identified by the Gene Meter approach were not identified by the DNA sequencing approach. The 52 microbial genera included: *Achromobacter, Actinobaculum,Afipia, Agrobacterium, Bartonella, Bdellovibrio, Brevundimonas, Bulleidia, Candida, Cardiobacterium, Caryophanon, Catonella, Centipeda, Chlamydia, Comamonas, Deinococcus, Delftia, Dermabacter, Desulfobulbus, Desulfomicrobium, Erythromicrobium, Eggerthella, Enterobacter, Escherichia, Ewingella, Flavobacterium, Flexistipes, Helicobacter, Holophaga, Janthinobacterium, Johnsonella, Klebsiella, Lautropia, Leptothrix, Leuconostoc, Methanobrevibacter, Mogibacterium, Ochrobactrum, Olsenella, Oribaculum, Pedobacter, Porphyromonas-like, Proteus, Rhizobium, Simonsiella, Solobacterium, Sphingomonas, Stenotrophomonas, Stomatococcus, Suttonella,* and *Tropheryma.*

It is important to note that the same genus (and species) could have different GI numbers. In the work below, the GI number was used (rather than the genus name) because there were instances when a genus with a certain GI was detected by both approaches but the same genus with a different GI number was not detected (details below). Altogether, the 52 genera were represented by 76 GI numbers. Since the average gene abundance maxima for these GIs was 8.6 a.u. (close to the maximum of 10 a.u.) in at least one patient, the microorganisms should have been identified in the DNA sequencing results. Putative reasons why they were not include: (i) the processing of the DNA sequencing reads, (ii) inadequate read depth, and (iii) false positive results, which are discussed below.

### Hidden jewels in the unprocessed sequencing reads

To identify microbial species, DNA sequencing reads are subjected to the various filtering procedures (e.g., singleton removal, minimum read lengths, similarities to existing rRNA databases, e.g., 97% similarity, 100 bp min alignment). The filtering can potentially remove 16S rRNA genes of microbial species that are actually present in the sample but not reported in the final taxonomic assignment. To demonstrate this phenomenon, one could match the unique probe sequences of each microbial species (not observed in the final taxonomic assignment) against all unfiltered DNA sequence reads (i.e., before taxonomic assignments are made). If any of the unique probes happen to match the unprocessed DNA sequences for a particular microbial species, one could infer that filtering procedures were responsible for not identifying the microorganism in the final DNA sequencing results. In other words, the rRNA genes of the microbial species were actually present in the unfiltered DNA sequence reads and indeed detected by the Gene Meter approach. However, they are not present in the final filtered DNA sequencing results.

Using custom-designed C++ programs and *Actinomyces* sp. (GI 2073386) as a positive control (since 16S rRNA gene of this microorganism was found in all patient samples), we matched the unique probes against the DNA sequencing reads of each patient sample. We found that 67 of the 76 GIs (∼88%) were present in the DNA sequencing reads. The 45 genera included: *Achromobacter, Actinobaculum, Agrobacterium, Bartonella, Bdellovibrio, Bulleidia, Candida, Cardiobacterium, Caryophanon, Catonella, Centipeda, Chlamydia, Deinococcus, Delftia, Dermabacter, Desulfobulbus, Desulfomicrobium, Eggerthella, Enterobacter, Erythromicrobium, Ewingella, Flavobacterium, Flexistipes, Helicobacter, Holophaga, Janthinobacterium, Johnsonella, Klebsiella, Lautropia, Leptothrix, Leuconostoc, Methanobrevibacter, Mogibacterium, Ochrobactrum, Olsenella, Oribaculum Pedobacter, Porphyromonas-like, Proteus, Simonsiella, Solobacterium, Sphingomonas, Stomatococcus, Suttonella,*and *Tropheryma.* A histogram of the frequencies of the unique probes for each of the 76 GIs revealed that most organisms had 50 or more unique probes matching to the DNA sequencing reads, with some having up to 400 matching probes (Figure 6).

**Figure 6.**
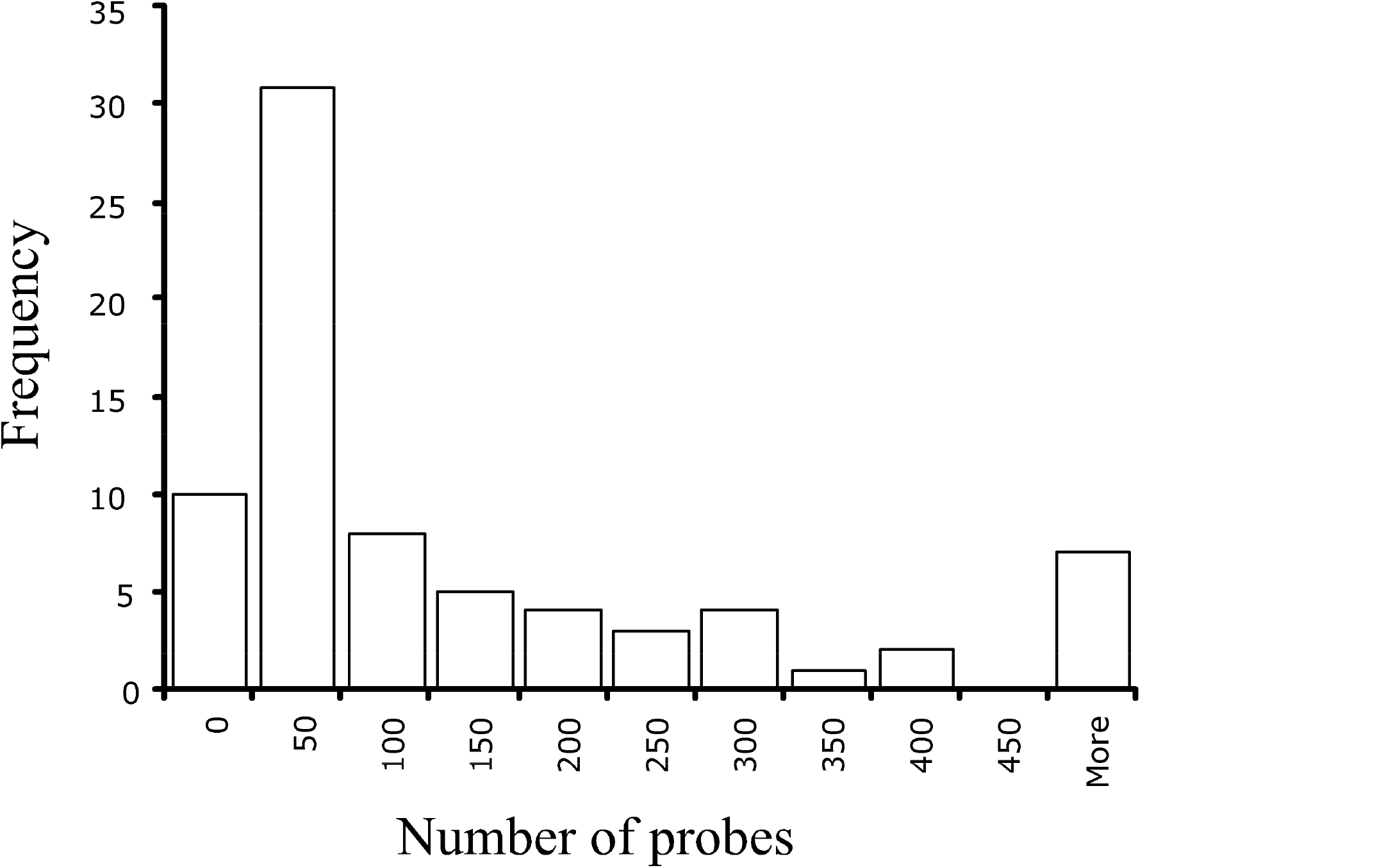
Number of probes of 76 microorganisms (GIs) belonging to the 52 genera (Figure 4) matching unprocessed DNA sequencing reads. Most microorganisms had 50 or more probe matches to sequencing reads, indicating they were in fact present in the patient samples (even though they were not detected by MG-RAST presumably due to filtering.

The significance of this result is that many microbial species – not taxonomically identified in DNA sequencing results – were actually present in the patient samples. Moreover, the inclusion of these results into the union of the two approaches increased the number of common microbial genera from 55 to 100 (Figure 4). In other words, 100 of the 107 genera (∼93%) identified by the Gene Meter approach were found in the processed and unprocessed DNA sequencing reads. These results indicate false negatives in the taxonomic assignment of the DNA sequencing results due to filtering and false positives in the Gene Meter results (since 7 were not identified in the sequencing reads).

### Microorganisms identified by the Gene Meter approach but not found in the unfiltered sequencing reads

Nine of the 76 GIs (∼12%) not detected in any of the unfiltered DNA sequencing reads included: *Afipia sp*. (*n*=191 probes)*, Brevundimonas diminuta* (*n*=224 probes)*, Caryophanon sp.* (*n*=53 probes)*, Comamonas sp.* (*n*=19 probes)*, Delftia sp.* (*n*=4 probes)*, Escherichia coli* (*n*=17 probes)*, Rhizobium loti* (*n*=335 probes), *Simonsiella muelleri* (*n*=31 probes), and *Stenotrophomonas maltophilia* (*n*=68 probes). The number of unique probes is shown because a microbial species with low number might not be detected in the unfiltered reads simply because of the low probability of finding a match. It should be noted that one species of *Leptothrix* (GI 14537943) with *n*=38 probes did not match any unfiltered sequence reads – however, another species of *Leptothrix* ( GI 14537937) with *n*=361 probes did match 96 sequencing reads. These findings indicate that the number of unique probes did influence whether or not a particular species was found in the unfiltered sequencing reads.

In our study, microorganisms with a low number of unique probes (*n*=38 or less) included: *Comamonas sp., Delftia sp., Simonsiella muelleri,* and *Escherichia coli*. We concluded that these microorganisms were probably not detected in the unfiltered sequencing reads because they had too few probes.

The remaining 5 species, detected by the Gene Meter approach but not detected in the unfiltered sequencing reads, had a high number of unique probes (i.e., *Afipia sp*.*, Brevundimonas diminuta, Caryophanon sp., Rhizobium loti,* and *Stenotrophomonas maltophilia)*. In theory, these microorganisms should have been detected in the unfiltered sequencing reads – but they were not. There are two possible reasons for this phenomenon. First, there might not have been sufficient read depth of the unfiltered sequencing data. According to our previous paper (Table S1 in Ref. 32), the read depth varied by patient sample with the lowest being 2,064 reads for a healthy patient and the highest being 23,904 reads for a caries patient. *Afipia sp.* for example, had high abundance in one of the caries patients (8.4 a.u.) and one of the patients with edentulism (7.6 a.u.) (determined by the Gene Meter approach) – yet, the read counts for these samples were 11,532 and 18,515, respectively, indicating this microorganism should have been identified in the unfiltered DNA sequencing reads. Hence, there is weak support for the argument since the highest abundances (by the Gene Meter approach) occurred in samples that also had moderate to high read counts.

Second, these microorganisms might not have been found in the unfiltered sequencing reads because they are false positives of the Gene Meter approach due to non-specific hybridization to other gene targets. We tested for the potential of non-specific hybridizations by removing the first three nucleotides on the 5’- and 3’-ends from the unique probes (because they might hybridize to non-specific DNA targets) and matching the short probes (19 nt) against the unprocessed sequencing reads. We found that several probes of *Caryophanon sp., and Rhizobium loti* and *Stenotrophomonas maltophilia* matched the unfiltered sequencing reads. Specifically, two unique probes (1015327 and 1015321) for *Caryophanon sp*. matched unfiltered sequence reads in a patient with caries and another patient with health. These probes have the potential to non-specifically bind to 16S rRNA sequences of other species in the patient samples and therefore they are false positives.

Ninety-six of the 151 genera (∼64%) identified by DNA sequencing were not detected by the Gene Meter approach included: *Acetitomaculum, Acetivibrio, Acetobacterium, Acidithiobacillus, Acidovorax, Actinoplanes, Aerococcus, Agrococcus, Alistipes, Aminobacterium, Amycolatopsis, Anaerostipes, Aquitalea, Arcanobacterium, Arthrobacter, Bavariicoccus, Bergeriella, Blautia, Brachymonas, Brenneria, Brevibacterium, Butyricimonas, Carnobacterium, Cellulophaga, Cellulosimicrobium, Chelonobacter, Chitinophaga, Chryseobacterium, Collinsella, Cryobacterium, Cytophaga, Dechloromonas, Desulfonispora, Desulfosporosinus, Desulfotomaculum, Enterorhabdus, Exiguobacterium, Finegoldia, Gallibacterium, Gardnerella, Geobacillus, Globicatella, Gordonibacte, Hespellia, Jonesia, Kocuria, Kytococcus, Lactococcus, Mannheimia, Megamonas, Melissococcus, Mobiluncus, Myroides, Odoribacter, Pantoea, Parabacteroides, Paraprevotella, Pasteurella, Pectinatus, Phascolarctobacterium, Promicromonospora, Pseudobutyrivibrio, Pseudonocardia, Pyramidobacter, Ralstonia, Renibacterium, Rhodococcus, Riemerella, Rikenella, Robinsoniella, Roseburia, Ruminococcus, Saccharopolyspora, Sanguibacter, Sebaldella, Serratia, Spirochaeta, Sporomusa, Streptobacillus, Streptomyces, Syntrophococcus, Tenacibaculum, Terrabacter, Tetragenococcus, Thermoanaerobacter, Thermomonospora, Thermus, Thiobacillus, Tissierella, Trichococcus, Vagococcus, Weeksella, Weissella,* and *Xylanimicrobium.* These microorganisms were not detected because the microarray probes did not target their 16S rRNA genes.

### An attempt to calibrate DNA sequencing data

We investigated the feasibility of calibrating the 454 sequencing reads by diluting pooled patient samples with DNA from salmon sperm. We chose salmon sperm because the sequences are dissimilar to microbial 16S rRNA genes and can easily be distinguished. In theory, DNA sequencing reads should yield similar linear equations for the dilution series but, in reality, amplification and sequencing biases vary for different targets and one would expect the slopes of the dilution series to be dissimilar (Figure 7). Our results show that the slope of *Neisseria* sp. was 69.0, while that of *Porphyromonas gingivalis* was 2818.2. Similarly, the slope of *Eubacterium brachy* was 102.7, while that of *Veillonella* sp. was 397.3. The importance of these findings are two-fold. First, it provides proof of the extreme biases of sequencing reads generated using NGS. Second, the results suggest that it might be possible to obtain precise abundances of 16S rRNA genes using NGS by calibrating 16S rRNA gene reads, before determining gene abundances, similar to what was done with the DNA microarray output using the Gene Meter approach. To our knowledge, this is the first study to provide preliminary evidence that gene abundances might be precisely determined by next-generation sequencing data.

**Figure 7.**
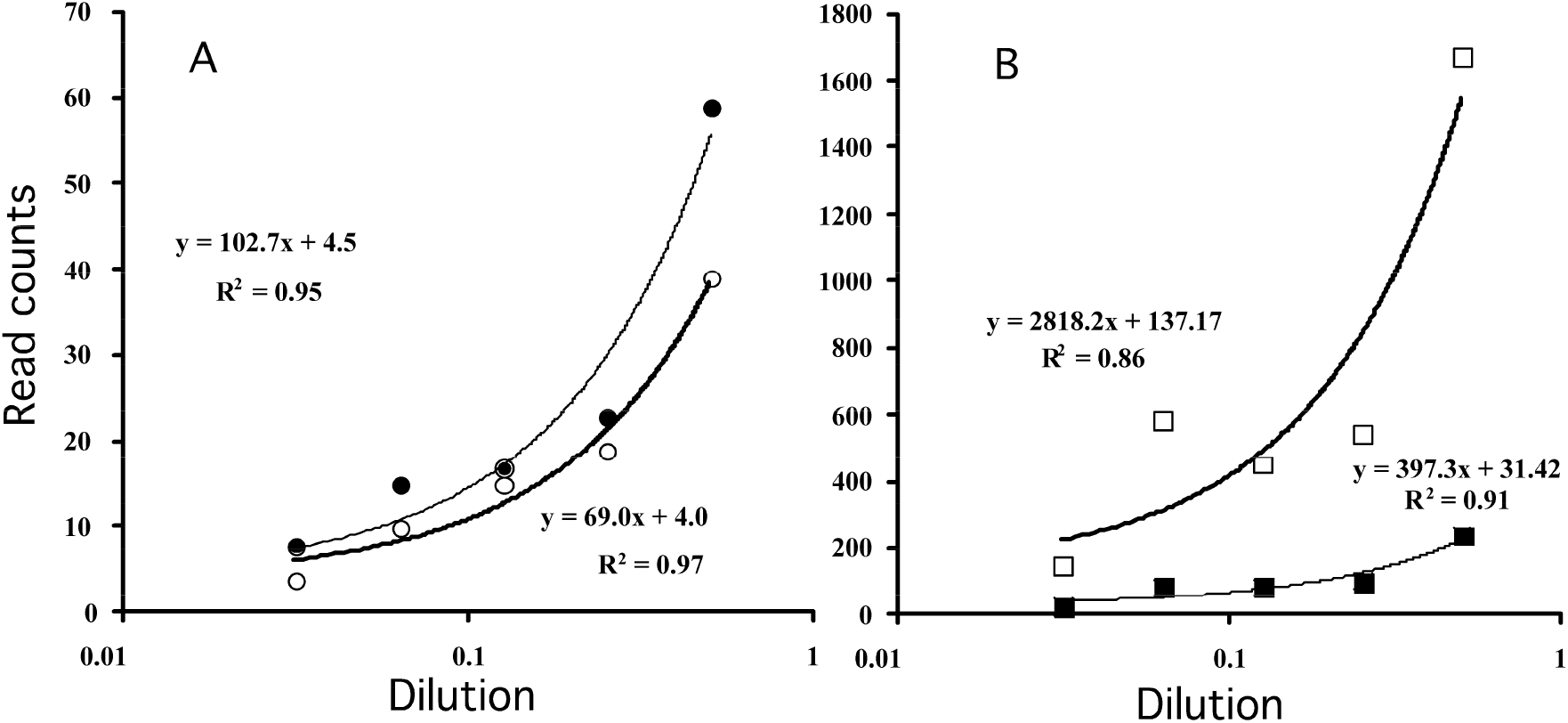
Calibration of 454 reads using the Gene Meter approach. Panel A: Neisseria sp., open circles; Eubacterium brachy, closed circles. Panel B: Porphyromonas gingivalis, open squares; Veillonella sp., closed squares.

## DISCUSSION

### DNA microarrays - too early to abandon

A common perception among researchers is that high-throughput DNA sequencing is the best approach to characterize microbial communities because other approaches, such as DNA microarrays are noisy. Indeed, conventional DNA microarrays are noisy (41) because the signal is not calibrated and the output has to be normalized for sample comparison. The Gene Meter approach solves the noise problem by calibrating all probes on a microarray before testing biological samples. The calibration identifies probes with noisy behavior because they will not fit the adsorption models. Noisy probes are not used in subsequent analyses. The Gene Meter approach also does not require any normalization of the output because the same amount of DNA is loaded onto every microarray. In other words, the abundances of different samples can be directly compared without introducing biases associated with other methods such as DNA sequencing and conventional DNA microarrays. It is for these reasons that we believe that the Gene Meter approach is superior to existing approaches – particularly when one desires to determine the abundance of a gene in a pool of genes.

It should be noted that DNA sequencing approaches (such as the one used in our previous study; 32) have problems too, that limit their ability to determine gene abundance. For example, an extra PCR amplification step is required in DNA sequencing (e.g., emulsion PCR) and that adds biases to the interpretation of the output. The Gene Meter approach does not need any extra amplification step – the labeled DNA is added directly to the microarray. Another problem in DNA sequencing (as shown in this study) is the post-processing of the reads sometimes filter out 16S rRNA genes that are actually present in biological samples (specifically, 40 out of 107 genera in this study). The fact that many of the unique probes of the 40 unidentified genera matched the unprocessed sequencing reads suggest that the taxonomic assignments in other sequencing studies might have grossly underestimated the actual diversity of microorganisms in biological samples.

We recognize that quantitative PCR (qPCR) could also have been used to determine the abundance of 16S rRNA genes in our study. However, like DNA sequencing, this approach is subject to PCR amplification biases and normalization steps. In addition, qPCR can only investigate a limited number of gene targets at one time (i.e., it is not high throughput), while the Gene Meter approach is high throughput but limited to the number of probes on microarray surface.

Similar to all approaches, the Gene Meter approach is not without its problems. For instance, seven genera identified by the Gene Meter approach were not detected in either the processed or unprocessed DNA sequencing reads. Three of the 7 genera were classified as false positives as they had the potential to cross-hybridize to the 16S rRNA genes of other species. Although there are several additional explanations for these phenomena (e.g., insufficient DNA sequencing depth; need for better probe design), the Gene Meter approach still provides precise target abundances for many genera and is superior to other approaches because normalization is not required.

A simple solution to prevent normalization biases and improve upon abundance determinations for DNA sequencing is to calibrate the instrument before analyzing samples. This study was the first to demonstrate that a DNA sequencing instrument can be calibrated using a dilution series (Figure 7). Future studies are now warranted to demonstrate the utility of calibration for DNA sequencing instruments using biological samples.

### Calibration by aggregate probes works optimally

In early Gene Meter studies (18,19), the dilution series were calibrated using the averaged signal intensities of all replicated probes specific to a gene target. The averaging of the signal reduced noise by minimizing the effects of outliers. Since we did not use identically replicated probes in the present study, we calibrated the DNA microarray two ways using individual probes or aggregated probes. We sought to determine which of the two calibrations was optimal.

Calibration of the individual probes involved modeling the signal intensities of an individual probe in a dilution series. The abundance of the gene in a biological sample was determined by back-calculating the abundances of all probes which target that specific gene using the calibration and then averaging (or determining the median) the gene abundances. For example, consider three calibrated probes targeting the same gene. The abundance of a specific gene by calibrated probe A was 1 a.u., probe B was 2 a.u. and probe C was 3 a.u. Therefore, the final abundance of the gene was the average or median, 2 a.u.

Calibration of the aggregated probes involved summing the signal intensities of all probes in a dilution series that target a specific gene and determining the model that best fits the experimental data. For example, if the signal intensities of probes A, B, and C for a specific gene were 300, 500, and 700 relative fluorescence units (RFU) at one dilution, then the signal intensity used to calibrate the gene would be 1500 RFU. The abundance of a gene in a biological sample was determined by summing the signal intensities of all probes that target the gene and back-calculating the relative abundance from the calibration.

What distinguishes the two calibration approaches is that the aggregate probe approach uses the sum of the signal intensities of all probes as input to the calibration model while the individual probe approach uses only the signal intensity of an individual probe as input to the calibration model. Also, in the aggregate probe approach, the abundance is directly determined from the model, while in the individual probe approach, the final gene abundance is determined by the averaged (or median) abundance of all probes targeting that gene.

We showed that calibrations of aggregated probes were better than individual probes when the gene target abundances were low. The reason for this phenomenon is that the low abundance genes are more affected by signal noise because some readings were close to the level of resolution of the microarray scanner. The sum of the signal intensities of all probes targeting a specific gene minimizes this problem because the noise is not averaged. Our results showed that if the gene target abundance was greater than 2 a.u. (Figure 4), there were no significant differences in the gene abundances by the calibration approach.

Interestingly, we found more differences in gene abundances by patient condition using aggregated probes (Tables 3, 4 and 5) than individual probes (Table 1 and 2). This finding suggests that several microorganisms do have differences in abundances by condition – but they occur at low abundances (<2 a.u.). In other words, the differences by condition could only be detected using the aggregated probe approach. Therefore, the aggregated probes was used for examining microbial species differences by condition (below).

### Abundance signatures for dysbiosis

The following 17 genera had significantly higher abundances in patients with 610 periodontitis and patients with edentulism when contrasted with patients with health: *Actinomyces, Bifidobacterium, Butyrivibrio, Campylobacter, Capnocytophaga, Eubacterium, Klebsiella, Lactobacillus, Leptotrichia, Neisseria, Peptostreptococcus, Porphyromonas, Prevotella, Selenomonas, Streptococcus, Treponema,* and *Veillonella.* At the species level this included: *Actinomyces odontolyticus, Klebsiella pneumoniae, Peptostreptococcus anaerobius, Streptococcus gordonii, Treponema socranskii,* and *Veillonella dispar.* We classified these microorganisms as ‘signatures of dysbiosis’ in the human oral microbiome because they occurred in more than one condition. They were excluded from consideration to a single condition below (e.g., a signature for periodontitis).

These microorganisms have previously been reported to be involved in dysbiosis in human oral studies. For example, *Actinomyces* sp. are found in edentulous patients (42). *Bifidobacterium* are found in the mouths of edentulous patient wearing dentures (43) and associated with root decay and periodontal disease (44). *Butyrivibrio, Campylobacter, Eubacterium, Prevotella, Selenomonas, Streptococcus, Leptotrichia,* and *Treponema* are associated with peri-implant communities (45). *Eubacterium, Prevotella* and *Lactobacillus* are major genera found in edentulous patients (46). *Prevotella, Fusobacterium*, *Leptotrichia, Streptococcus, Neisseria,* and *Veillonella* were found in edentulous infants (24,47,48). *Peptostreptococcus* has been associated with periodontal pockets of partially edentate patients (49) and is associated with soft tissue inflammation in edentate patients (42). *Prevotella* and *Campylobacter* have been observed in edentate patients (42). *Treponema Campylobacter, Peptostreptococcus,* and *Porphyromonas* are associated with periodontitis (50, 51). *Veillonella, Streptococcus* and *Candida* are often found in the saliva of edentulous patients (52).

At the species level, *Actinomyces odontolyticus* has been found in failed dental implants (53). *Klebsiella pneumoniae* has been frequently observed in patients with diseased implants (54) and patients with periodontitis (55,56). *Peptostreptococcus anaerobius* has been found inside root canals (50). *Streptococcus gordonii* has been shown to be an important player in the development of periodontitis as it is involved in co-adhesion and metabolic interactions with other pathogens (53,57). *Treponema socranskii* has been associated with peri-implant diseases (58,59), lesions of teeth with apical periodontitis (60), infected root canals and abscesses (61). *Veillonella dispar* is an earlier biofilm colonizer (62) and involved in periodontal disease (63).

### Abundance signatures for periodontitis

The following 13 genera had significantly higher abundances in patients with periodontitis than those with health: *Abiotrophia, Bacteroides-like, Caryophanon, Clostridium, Dialister, Filifactor, Fusobacterium, Granulicatella, Haemophilus, Megasphaera, Mogibacterium, Staphylococcus,* and *Tannerella.* At the species level this included: *A. elegans, C. tetanomorphum, F. alocis, F. nucleatum, G. elegans, H. segnis, M. diversum, S. epidermidis, S. xylosus,* and *T. forsythia.* Note that some genera could not be resolved to the species level. These microorganisms were classified as microbial signatures of periodontitis in the oral microbiome.

The following microorganisms have been previously shown to be associated with periodontal disease: *Abiotrophia elegans* (64,65), *Bacteroides-like* (66), *Dialister* (67,68,69), *Filifactor alocis* (70,71,72,73,74), *Fusobacterium nucleatum* (15,42,75,76,77,78), *Granulicatella elegans* (79), *Haemophilus* (70,80,81), *Haemophilus segnis* (82,83) *Megasphaera* (4,84), *Mogibacterium* (85,86), *Staphylococcus epidermidis* (87), *Tannerella forsythia* (71,77,89,90). These findings support our findings that these microorganisms are signatures of periodontitis in the human oral microbiome.

Although the genera *Clostridium Mogibacterium* and *Staphylococcus* have been associated with periodontal disease (10,91), there is no evidence in the literature suggesting that *Clostridium tetanomorphum, Mogibacterium diversum, Staphylococcus xylosus* or the genus *Caryophanon* has been associated with periodontitis. Of note, *Mogibacterium diversum* has been associated with caries (88) and the genera *Caryophanon* has been found in the gingival flora of dogs (92). Recall that our earlier results suggest that *Caryophanon* might be a false positive because it was not found in the sequencing reads and some probes were shown to have the potential to hybridize with other species.

A probable reason these microorganisms were not recognized in the literature is because they occurred at low abundances and might be overlooked using alternative technologies (e.g., NGS). For example, of the 525 *Clostridium tetanomorphum* probes, the abundance in patients with health was 1.8 ± 0.50 a.u. versus 3.1 ± 0.81 a.u. in patients with periodontitis. Similarly, of the 114 *Mogibacterium diversum* probes, the abundance in patients with health was 1.6 ± 0.30 a.u. versus 2.8 ± 0.72 a.u. in patients with periodontitis, and of the 54 *Staphylococcus xylosus* probes, the abundance in patients with health was 1.5 ± 0.25 a.u. versus 2.4 ± 0.48 a.u. in patients with periodontitis (Table 5). All comparisons are statistically significant and all species have been previously found in human oral cavities.

### Abundance signatures for edentulism

The following 14 genera had significantly higher abundances in patients with edentulism than patients with health: *Atopobium, Bacillus, Burkholderia, Candida, Deinococcus, Erysipelothrix, Ewingella, Kingella, Mycoplasma, Paenibacillus, Pedobacter, Propionibacterium, Solobacterium,* and *Stomatococcus.* At the species level, this included: *Atopobium rimae, Bacillus clausii, Candida albicans, Erysipelothrix tonsillarum, Ewingella americana, Kingella denitrificans, Mycoplasma buccale, Propionibacterium avidum,* and *Stomatococcus mucilaginosus.* While high abundances of *Atopobium* species, *Candida albicans* and *Bacillus sp.* have been previously reported in edentulous patients (93), we could not find any evidence for the high abundance of the other genera and species in patients with edentulism in the literature. Hence, this is the first study to show higher abundance of these microorganisms in patients with edentulism.

### Technology rules our understanding of the oral microbiome

Much of what is known about the oral microbiome is based on the technology available at the time of publication. For example, the idea behind the ‘red complex’ originated from the checkerboard DNA–DNA hybridization technique, which was used to examine the microbial composition of plaque in patients in health and periodontitis (94,95), the salivary microbiota levels in relation to periodontal status (96), the relationship of cigarette smoking to the composition of the subgingival microbiota (97,98), the differences between the subgingival microbiota in patients from different geographic locations (99), the relationship of ethnic/racial group, occupational and periodontal disease status (100), and effects of different periodontal therapies (101,102). While the technique is rapid, sensitive, and relatively inexpensive, its major shortcoming was cross-hybridization (103). The implementation of new technologies such as DNA microarrays and next-generation-sequencing improved upon our understanding of the human oral microbiome (28) – but these technologies are not really quantitative. Gene Meter methodology allows researcher to precisely determine microbial abundances, which allows for statistical comparisons of different oral clinical conditions. We used these comparisons to find abundance signatures for dysbiosis, periodontitis and edentulism. These signatures could be used, individually or in combination, to assess the clinical status of a patient (e.g., 697 evaluating treatments such as antibiotic therapies, and oral microbial transplantations, e.g., 105).

## CONTRIBUTIONS

Experimental design: PAN and AP.

Laboratory Work: MCH, PAN, and AP.

Bioinformatic Work: PAN and AP.

Manuscript writing: MCH, PAN, and AP.

## Supplementary Materials

Table S1. GI numbers for 16S rRNA gene sequences of 597 oral bacteria.

Table S2. Median abundances for the 576 rRNA genes and 16 patient samples based on individual probe calibration.

Table S3. Abundances for the 567 rRNA genes and 16 patient samples based on aggregate probe calibration.

## LITERATURE CITED

1. Armitage GC. Development of a classification system for periodontal diseases and conditions. Ann Periodontol. 1999; 4:1–6. PMID: 10863370

2. Dewhirst FE, Chen T, Izard J, Paster BJ, Tanner AC, Yu WH, Lakshmanan A, Wade WG. The human oral microbiome. J Bacteriol. 2010; 192:5002–17. doi:10.1128/JB.00542-10. PMID: 20656903

3. Holt SC, Ebersole JL. Porphyromonas gingivalis, Treponema denticola, and Tannerella forsythia: the ``red complex’’, a prototype polybacterial pathogenic consortium in periodontitis. Periodontol 2000., 2005; 38:72–122. PMID: 15853938

4. Kumar PS, Griffen AL, Barton JA, Paster BJ, Moeschberger ML, Leys EJ. New bacterial species associated with chronic periodontitis. J Dent Res. 2003; 82:338–44. PMID: 12709498

5. Darveau RP. Periodontitis: a polymicrobial disruption of host homeostasis. Nat Rev Microbiol. 2010; 8:481–90. doi:10.1038/nrmicro2337. PMID: 20514045

6. Hajishengallis G, Liang S, Payne MA, Hashim A, Jotwani R, Eskan MA, McIntosh ML, Alsam A, Kirkwood KL, Lambris JD, Darveau RP, Curtis MA. Low-abundance biofilm species orchestrates inflammatory periodontal disease through the commensal microbiota and complement. Cell Host Microbe. 2011; 10:497–506. doi:10.1016/j.chom.2011.10.006. PMID: 22036469

7. Diaz PI, Dupuy AK, Abusleme L, Reese B, Obergfell C, Choquette L, Dongari-Bagtzoglou A, Peterson DE, Terzi E, Strausbaugh LD. Using high throughput sequencing to explore the biodiversity in oral bacterial communities. Mol Oral Microbiol. 2012; 27:182–201. doi:10.1111/j.2041-1014.2012.00642.x. PMID: 22520388

8. Bik EM, Long CD, Armitage GC, Loomer P, Emerson J, Mongodin EF, Nelson KE, Gill SR, Fraser-Liggett CM, Relman DA. Bacterial diversity in the oral cavity of 10 healthy individuals. ISME J. 2010; 4:962–74. doi:10.1038/ismej.2010.30. PMID: 20336157

9. Liu B, Faller LL, Klitgord N, Mazumdar V, Ghodsi M, Sommer DD, Gibbons TR, Treangen TJ, Chang YC, Li S, Stine OC, Hasturk H, Kasif S, Segrè D, Pop M, Amar S. Deep sequencing of the oral microbiome reveals signatures of periodontal disease. PLoS One. 2012; 7:e37919. doi:10.1371/journal.pone.0037919. PMID: 22675498

10. Griffen AL, Beall CJ, Campbell JH, Firestone ND, Kumar PS, Yang ZK, Podar M, Leys EJ. Distinct and complex bacterial profiles in human periodontitis and health revealed by 16S pyrosequencing. ISME J. 2012; 6:1176–1185. PMID: 22170420

11. Zaura E, Keijser BJ, Huse SM, Crielaard W. Defining the healthy ``core microbiome’’ of oral microbial communities. BMC Microbiol. 2009; 15:259. PMID: 22313693

12. Hajishengallis G, Lamont RJ. Beyond the red complex and into more complexity: the polymicrobial synergy and dysbiosis (PSD) model of periodontal disease etiology. Molecul Oral Microbiol, 2012; 27:409–419. PMID: 23134607

13. Huyghe A, Francois P, Charbonnier Y, Tangomo-Bento M, Bonetti EJ, Paster BJ, Bolivar I, Baratti-Mayer D, Pittet D, Schrenzel J; Geneva Study Group on Noma (GESNOMA). Novel microarray design strategy to study complex bacterial communities. Appl Environ Microbiol. 2008; 74:1876–85. doi:10.1128/AEM.01722-07. PMID: 18203854

14. Iwai S, Fei M, Huang D, Fong S, Subramanian A, Grieco K, Lynch SV, Huang L. Oral and airway microbiota in HIV-infected pneumonia patients. J Clin Microbiol. 2012; 50:2995–3002. doi:10.1128/JCM.20198-12. PMID: 22760045

15. Topcuoglu N, Kulekci G. 16S rRNA based microarray analysis of ten periodontal bacteria in patients with different forms of periodontitis. Anaerobe. 2015; 35(Pt A):35–40. doi:10.1016/j.anaerobe.2015.01.011. PMID: 25638399

16. Roth SB, Jalava J, Ruuskanen O, Ruohola A, Nikkari S. Use of an oligonucleotide array for laboratory diagnosis of bacteria responsible for acute upper respiratory infections. J Clin Microbiol. 2004; 42:4268–74. PMID: 15365022

17. Pozhitkov AE, Bailey KD, Noble PA. Development of a statistically robust quantification method for microorganisms in mixtures using oligonucleotide microarrays. J Microbiol Methods. 2007; 70:292–300. PMID: 17553581

18. Pozhitkov A.E., P.A. Noble, J. Bryk, D. Tautz. A revised design for microarray experiments to account for experimental noise and the uncertainty of probe response. Plos One 2014; 9:e91295 PMID: 24618910

19. Harrison A., Binder, H. Buhot, A., Burden, C.J., Carlon, E., Gibas, C., Gamble, L.J. Halperin, A., Hooyberghs, J., Kreil, D.P., Levicky, R., Noble, P.A., Ott, A., Pettitt, B.M., Tautz, D., Pozhitkov, A.E. Physico-chemical foundations underpinning microarray and next-generation sequencing experiments. Nucl Acids Res. 2013; 41:2779–96. PMID: 23307556

20. Hunter M.C., Pozhitkov A.E. and Noble P.A. Accurate predictions of postmortem interval using linear regression analyses of gene meter expression data. 2016; http://www.biorxiv.org/content/early/2016/06/12/058370

21. Pozhitkov A.E., Neme R., Domazet-Loo T., Leroux B.G., Soni S., Tautz D., and Noble P.A. Thanatotranscriptome: genes actively expressed after organismal death. 2016; http://www.biorxiv.org/content/early/2016/06/11/058305

22. Nasidze I, Li J, Quinque D, Tang K, Stoneking M. Global diversity in the human salivary microbiome. Genome Res. 2009; 19:636–43. doi:10.1101/gr.084616.108. Epub 2009 Feb 27. PMID: 19251737

23. Ahn J, Yang L, Paster BJ, Ganly I, Morris L, Pei Z, Hayes RB. Oral microbiome profiles: 16S rRNA pyrosequencing and microarray assay comparison. PLoS One. 2011; 6:e22788. doi:10.1371/journal.pone.0022788. PMID: 21829515

24. Cephas KD, Kim J, Mathai RA, Barry KA, Dowd SE, Meline BS, Swanson KS. Comparative analysis of salivary bacterial microbiome diversity in edentulous infants and their mothers or primary care givers using pyrosequencing. PLoS One. 2011; 6:e23503. doi:10.1371/journal.pone.0023503. PMID: 21853142

25. Simón-Soro A, Tomás I, Cabrera Rubio R, Catalan MD, Nyvad B, Mira A. Microbial geography of the oral cavity. J Dent Res. 2013; 92:616–21. doi:10.1177/0022034513488119. PMID: 23674263

26. Xu H, Hao W, Zhou Q, Wang W, Xia Z, Liu C, Chen X, Qin M, Chen F. Plaque bacterial microbiome diversity in children younger than 30 months with or without caries prior to eruption of second primary molars. PLoS One. 2014; 9:e89269. doi:10.1371/journal.pone.0089269. PMID: 24586647

27. Sato Y, Yamagishi J, Yamashita R, Shinozaki N, Ye B, Yamada T, Yamamoto M, Nagasaki M, Tsuboi A. Inter-individual differences in the oral bacteriome are greater than intra-day fluctuations in individuals. PLoS One. 2015; 10:e0131607. doi:10.1371/journal.pone.0131607. PMID: 26121551

28. Pozhitkov AE, Beikler T, Flemmig T, Noble PA. High-throughput methods for analysis of the human oral microbiome. Periodontol 2000. 2011;55:70–86. doi:10.1111/j.1600-0757.2010.00380.x. PMID: 21134229

29. Amend Anthony S, Seifert Keith A., Bruns Thomas D. Quantifying microbial communities with 454 pyrosequencing: does read abundance count? Molecular Ecol. 2010; 19:I24. PMID: 21050295

30. Dannemiller, N. Lang-Yona, N. Yamamoto, Y. Rudich, J. Peccia. Combining real-time PCR and next-generation DNA sequencing to provide quantitative comparisons of fungal aerosol populations. Atmospheric Environment, 2014; 84:113–121.

31. Jiang L, Schlesinger F, Davis CA, Zhang Y, Li R, Salit M, Gingeras TR, Oliver B. Synthetic spike in standards for RNA-seq experiments. Genome Res. 2011; 21:1543–51. PMID: 21816910

32. Pozhitkov A.E., Leroux B., Randolph T.W., Beikler T., Flemmig T.F., and Noble P.A. Towards microbiome transplant as a therapy for periodontitis: an exploratory study of periodontitis microbial signature contrasted by oral health, caries and edentulism. BMC Oral Health 2015; 15:125 PMID: 26468081

33. Page RC, Eke PI. Case definitions for use in population-based surveillance of periodontitis. J Periodontol. 2007; 78:1387–99. PMID: 17608611

34. Dye BA, Tan S, Smith V, Lewis BG, Barker LK, Thornton-Evans G, et al. Trends in oral health status: United States, 1988–1994 and 1999–2004. National Center for Health Statistics. Vital Health Stat. 2007; 11:248.

35. Beikler T, Schnitzer S, Abdeen G, Ehmke B, Eisenacher M, Flemmig TF. Sampling strategy for intraoral detection of periodontal pathogens before and following periodontal therapy. J Periodontol. 2006; 77:1323–3132. PMID: 16881801

36. Flemmig TF, Arushanov D, Daubert D, Rothen M, Mueller G, Leroux BG. Randomized controlled trial assessing efficacy and safety of glycine powder air polishing in moderate to deep periodontal pockets. J Periodontol. 2012; 83:444–52. PMID: 21861637

37. Meyer F, Paarmann D, D’Souza M, Olson R, Glass EM, Kubal M, Paczian T, Rodriguez A, Stevens R, Wilke A, Wilkening J, Edwards RA. The metagenomics RAST server - a public resource for the automatic phylogenetic and functional analysis of metagenomes. BMC Bioinf. 2008; 9:386. PMID: 18803844

38. Griffen AL, Beall CJ, Firestone ND, Gross EL, Difranco JM, Hardman JH, Vriesendorp B, Faust RA, Janies DA, Leys EJ. CORE: a phylogenetically-curated 16S rDNA database of the core oral microbiome. PLoS One. 2011. Apr 22;6(4):e19051. doi:10.1371/journal.pone.0019051. PMID: 21544197

39. Pozhitkov A, Noble PA, Domazet-Loso T, Nolte AW, Sonnenberg R, Staehler P, Beier M, Tautz D. Tests of rRNA hybridization to microarrays suggest that hybridization characteristics of oligonucleotide probes for species discrimination cannot be predicted. Nucleic Acids Res. 2006; 1734:e66. PMID: 16707658

40. Diaz PI, Chalmers NI, Rickard AH, Kong C, Milburn CL, Palmer RJ Jr, Kolenbrander PE. Molecular characterization of subject-specific oral microflora during initial colonization of enamel. Appl Environ Microbiol. 2006; 72:2837–2848. PMID: 16597990

41. Tu Y,Stolovitzky G, Klein U. Quantitative noise analysis for gene expression microarray experiments. Proc Natl Acad Sci USA. 2002; 99:14031. PMID: 12388780

42. Danser MM, vanWinkelhoff AJ, van der Velden U. Periodontal 841 bacteria colonizing oral mucous membranes in edentulous patients wearing dental implants. J Periodontol. 1997; 68:209–16.

43. Mantzourani M, Gilbert SC, Fenlon M, Beighton D. Non-oral bifidobacteria and the aciduric microbiota of the denture plaque biofilm. Mol Oral Microbiol. 2010; 25:190–9. doi:10.1111/j.2041-1014.2009.00565.x. PMID: 20536746

44. Dina MN, Mărgărit R, Andrei OC. Pontic morphology as local risk factor in root decay and periodontal disease. Rom J Morphol Embryol. 2013; 54:361–4. PMID: 23771082

45. Kumar PS, Mason MR, Brooker MR, O’Brien K. Pyrosequencing reveals unique microbial signatures associated with healthy and failing dental implants. J Clin Periodontol. 2012; 39:425–33. doi:10.1111/j.1600-051X.2012.01856.x. PMID: 22417294

46. Könönen E, Asikainen S, Alaluusua S, Könönen M, Summanen P, Kanervo A, Jousimies-Somer H. Are certain oral pathogens part of normal oral flora in denture-wearing edentulous subjects? Oral Microbiol Immunol. 1991; 6:119–22. PMID: 1945487

47. Teles FR, Teles RP, Sachdeo A, Uzel NG, Song XQ, Torresyap G, Singh M, Papas A, Haffajee AD, Socransky SS. Comparison of microbial changes in early redeveloping biofilms on natural teeth and dentures. J Periodontol. 2012; 83:1139–48. doi:10.1902/jop.2012.110506. PMID: 22443543

48. Könönen E, Asikainen S, Jousimies-Somer H. The early colonization of gram-negative anaerobic bacteria in edentulous infants. Oral Microbiol Immunol. 1992; 7:28–31. PMID: 1528621

49. van Winkelhoff AJ, Loos BG, van der Reijden WA, van der Velden U. Porphyromonas gingivalis, Bacteroides forsythus and other putative periodontal pathogens in subjects with and without periodontal destruction. J Clin Periodontol. 2002; 29:1023–8. PMID: 12472995

50. Piovano, S. Bacteriology of most frequent oral anaerobic infections. Anaerobe 1999; 5:221–227

51. van Winkelhoff AJ, Goené RJ, Benschop C, Folmer T. Early colonization of dental implants by putative periodontal pathogens in partially edentulous patients. Clin Oral Implants Res. 2000; 11:511–20. PMID: 11168244

52. Sato M, Hoshino E, Nomura, S, Ishioka K. Salivary microflora of geriatric edentulous persons wearing dentures. Microbial ecology in health and disease. 1993; 6:293–299.

53. Sakanaka A, Kuboniwa M, Takeuchi H, Hashino E, Amano A. Arginine-ornithine antiporter ArcD controls arginine metabolism and interspecies biofilm development of Streptococcus gordonii. J Biol Chem. 2015; 290:21185–98. doi:10.1074/jbc.M115.644401. PMID: 26085091

54. Sarkonen N, Könönen E, Eerola E, Könönen M, Jousimies-Somer H, Laine P. Characterization of Actinomyces species isolated from failed dental implant fixtures. Anaerobe. 2005;11:231–7. PMID: 16701573

55. Ardila CM, Fernández N, Guzmán IC. Antimicrobial susceptibility of moxifloxacin against gram negative enteric rods from Colombian patients with chronic periodontitis. J Periodontol. 2010; 81:292–9. doi:10.1902/jop.2009.090464. PMID: 20151809

56. Gonçalves MO, Coutinho-Filho WP, Pimenta FP, Pereira GA, Pereira JA, Mattos-Guaraldi AL, Hirata R Jr. Periodontal disease as reservoir for multi-resistant and hydrolytic enterobacterial species. Lett Appl Microbiol. 2007;44:488–94. PMID: 17451514

57. Hajishengallis G, Lamont RJ. Dancing with the stars: how choreographed bacterial interactions dictate nososymbiocity and give rise to keystone pathogens, accessory pathogens, and pathobionts. Trends Microbiol. 2016; 24:477–89. doi:10.1016/j.tim.2016.02.010.

58. Máximo MB, de Mendonça AC, Renata Santos V, Figueiredo LC, Feres M, Duarte PM. Short-term clinical and microbiological evaluations of peri-implant diseases before and after mechanical anti-infective therapies. Clin Oral Implants Res. 2009; 20:99–108. doi:10.1111/j.1600-0501.2008.01618.x. PMID: 19126114

59. Persson GR, Renvert S. Cluster of bacteria associated with peri-implantitis. Clin Implant Dent Relat Res. 2014;16:783–93. doi:10.1111/cid.12052. PMID: 23527870

60. Rosa TP, Signoretti FG, Montagner F, Gomes BP, Jacinto RC. Prevalence of Treponema spp. in endodontic retreatment-resistant periapical lesions. Braz Oral Res. 2015;29. pii: S1806-83242015000100228. doi:10.1590/1807-3107BOR-2015.vol29.0031. PMID: 25627883

61. Montagner F, Jacinto RC, Signoretti FG, Gomes BP. Treponema species detected in infected root canals and acute apical abscess exudates. J Endod. 2010;36:1796–9. doi:10.1016/j.joen.2010.08.008. PMID: 20951290

62. Mashima I, Nakazawa F. The influence of oral Veillonella species on biofilms formed by Streptococcus species. Anaerobe. 2014; 28:54–61. doi:10.1016/j.anaerobe.2014.05.003. PMID: 24862495

63. Moon JH, Lee JH, Lee JY. Subgingival microbiome in smokers and non-smokers in Korean chronic periodontitis patients. Mol Oral Microbiol. 2015;30:227–41. doi:10.1111/omi.12086. PMID: 25283067

64. Rego RO, Oliveira CA, dos Santos-Pinto A, Jordan SF, Zambon JJ, Cirelli JA, Haraszthy V. Clinical and microbiological studies of children and adolescents receiving orthodontic treatment. Am J Dent. 2010. Dec;23(6):317–23. PMID: 21344829

65. Mikkelsen L, Theilade E, Poulsen K. Abiotrophia species in early dental plaque. Oral Microbiol Immunol. 2000;15:263–8. PMID: 11154413

66. Olsvik B, Flynn MJ, Tenover FC, Slots J, Olsen I. Tetracycline resistance in Prevotella isolates from periodontally diseased patients is due to the tet(Q) gene. Oral Microbiol Immunol. 1996; 11:304–8. PMID: 9028255

67. Lee MY, Kim YJ, Gu HJ, Lee HJ. A case of bacteremia caused by Dialister pneumosintes and Slackia exigua in a patient with periapical abscess. Anaerobe. 2016; 38:36–8. doi:10.1016/j.anaerobe.2015.11.006. PMID: 26612007

68. Gonçalves C, Soares GM, Faveri M, Pérez-Chaparro PJ, Lobão E, Figueiredo LC, Baccelli GT, Feres M. Association of three putative periodontal pathogens with chronic periodontitis in Brazilian subjects. J Appl Oral Sci. 2016; 24:181–5. doi:10.1590/1678-775720150445. PMID: 27119767

69. Collins JR, Arredondo A, Roa A, Valdez Y, León R, Blanc V. Periodontal pathogens and tetracycline resistance genes in subgingival biofilm of periodontally healthy and diseased Dominican adults. Clin Oral Investig. 2016; 20:349–56. doi:10.1007/s00784-015-1516-2. PMID: 26121972

70. Colombo AP, Boches SK, Cotton SL, Goodson JM, Kent R, Haffajee AD, Socransky SS, Hasturk H, Van Dyke TE, Dewhirst F, Paster BJ. Comparisons of subgingival microbial profiles of refractory periodontitis, severe periodontitis, and periodontal health using the human oral microbe identification microarray. J Periodontol. 2009; 80:1421–32. doi:10.1902/jop.2009.090185. PMID: 19722792

71. Oliveira RR, Fermiano D, Feres M, Figueiredo LC, Teles FR, Soares GM, Faveri M. Levels of candidate periodontal pathogens in subgingival biofilm. J Dent Res. 2016; 95:711–8. doi:10.1177/0022034516634619. PMID: 26936213

72. Basic A, Dahlén G. Hydrogen sulfide production from subgingival plaque samples. Anaerobe. 2015; 35(Pt A):21–7. doi:10.1016/j.anaerobe.2014.09.017. PMID: 25280920

73. Aruni AW, Mishra A, Dou Y, Chioma O, Hamilton BN, Fletcher HM. Filifactor alocis–a new emerging periodontal pathogen. Microbes Infect. 2015; 17:517–30. doi:10.1016/j.micinf.2015.03.011. PMID: 25841800

74. Chen H, Liu Y, Zhang M, Wang G, Qi Z, Bridgewater L, Zhao L, Tang Z, Pang X. A Filifactor alocis- centered co-occurrence group associates with periodontitis across different oral habitats. Sci Rep. 2015; 5:9053. doi:10.1038/srep09053. PMID: 25761675

75. Ahn SH, Song JE, Kim S, Cho SH, Lim YK, Kook JK, Kook MS, Lee TH. NOX1/2 activation in human gingival fibroblasts by Fusobacterium nucleatum facilitates attachment of Porphyromonas gingivalis. Arch Microbiol. 2016; 198:573–83. doi:10.1007/s00203-016-1223-7. PMID: 27071620

76. Mendes RT, Nguyen D, Stephens D, Pamuk F, Fernandes D, Van Dyke TE, Kantarci A. Endothelial cell response to Fusobacterium nucleatum. Infect Immun. 2016; 84:2141–8. doi:10.1128/IAI.01305-15. PMID: 27185790

77. Nickles K, Scharf S, Röllke L, Mayer I, Mayer M, Eickholz P. Detection of subgingival periodontal pathogens–comparison of two sampling strategies. Clin Oral Investig. 2016; 20:571–9. doi:10.1007/s00784-015-1530-4. PMID: 26193958

78. Bui FQ, Johnson L, Roberts J, Hung SC, Lee J, Atanasova KR, Huang PR, Yilmaz Ö3 Ojcius DM. Fusobacterium nucleatum infection of gingival epithelial cells leads to NLRP3 inflammasome-dependent secretion of IL-1β and the danger signals ASC and HMGB1. Cell Microbiol. 2016; 18:970–81. doi:10.1111/cmi.12560. PMID: 26687842

79. Lourenço TG, Heller D, Silva-Boghossian CM, Cotton SL, Paster BJ, Colombo AP. Microbial signature profiles of periodontally healthy and diseased patients. J Clin Periodontol. 2014; 41:1027–36. doi:10.1111/jcpe.12302. PMID: 25139407

80. Park OJ, Yi H, Jeon JH, Kang SS, Koo KT, Kum KY, Chun J, Yun CH, Han SH. Pyrosequencing Analysis of Subgingival Microbiota in Distinct Periodontal Conditions. J Dent Res. 2015; 94:921–7. doi:10.1177/0022034515583531. PMID: 25904141

81. Takeshita T, Matsuo K, Furuta M, Shibata Y, Fukami K, Shimazaki Y, Akifusa S, Han DH, Kim HD, Yokoyama T, Ninomiya T, Kiyohara Y, Yamashita Y. Distinct composition of the oral indigenous microbiota in South Korean and Japanese adults. Sci Rep. 2014; 4:6990. doi:10.1038/srep06990. PMID: 25384884

82. Petsios A, Nakou M, Manti F. Microflora in adult periodontitis. J Periodontal Res. 1995; 30:325–31. PMID: 7494174

83. Kamma JJ, Nakou M, Manti FA. Predominant microflora of severe, moderate and minimal periodontal lesions in young adults with rapidly progressive periodontitis. J Periodontal Res. 1995; 30:66–72. PMID: 7722848

84. Kumar PS, Griffen AL, Moeschberger ML, Leys EJ. Identification of candidate periodontal pathogens and beneficial species by quantitative 16S clonal analysis. J Clin Microbiol. 2005; 43:3944–55. PMID: 16081935

85. Marchesan JT, Morelli T, Moss K, Barros SP, Ward M, Jenkins W, Aspiras MB, Offenbacher S. Association of Synergistetes and Cyclodipeptides with periodontitis. J Dent Res. 2015; 94:1425–31. doi:10.1177/0022034515594779. PMID: 26198391

86. Camelo-Castillo AJ, Mira A, Pico A, Nibali L, Henderson B, Donos N, Tomás I. Subgingival microbiota in health compared to periodontitis and the influence of smoking. Front Microbiol. 2015; 6:119. doi:10.3389/fmicb.2015.00119. PMID: 25814980

87. Fujii R, Saito Y, Tokura Y, Nakagawa KI, Okuda K, Ishihara K. Characterization of bacterial flora in persistent apical periodontitis lesions. Oral Microbiol Immunol. 2009; 24:502–5. doi:10.1111/j.1399-302X.2009.00534.x. PMID: 19832803

88. Nakazawa F, Poco SE Jr, Sato M, Ikeda T, Kalfas S, Sundqvist G, Hoshino E. Taxonomic characterization of Mogibacterium diversum sp. nov. and Mogibacterium neglectum sp. nov., isolated from human oral cavities. Int J Syst Evol Microbiol. 2002; 52(Pt 1):115–22. PMID: 11837293

89. Wong BK, McGregor NR, Butt HL, Knight R, Liu LY, Darby IB. Association of clinical parameters with periodontal bacterial haemolytic activity. J Clin Periodontol. 2016; 43:503–11. doi:10.1111/jcpe.12554. PMID: 27105613

90. Lanza E, Magan-Fernandez A, Bermejo B, de Rojas J, Marfil-Alvarez R, Mesa F. Complementary clinical effects of red complex bacteria on generalized periodontitis in a caucasian population. Oral Dis. 2016; 22:430–7. doi:10.1111/odi.12471. PMID: 26948988

91. VieiraColombo AP, Magalhães CB, Hartenbach FA, Martins do Souto R, Maciel da Silva-Boghossian C. Periodontal-disease-associated biofilm: A reservoir for pathogens of medical importance. Microb Pathog. 2016; 94:27–34. doi:10.1016/j.micpath.2015.09.009. PMID: 26416306

92. Nyby MD, Gregory DA, Kuhn DA, Pangborn J. Incidence of Simonsiella in the oral cavity of dogs. J Clin Microbiol. 1977; 6:87–8. PMID: 886011

93. O’Donnell LE, Robertson D, Nile CJ, Cross LJ, Riggio M, Sherriff A, Bradshaw D, Lambert M, Malcolm J, Buijs MJ, Zaura E, Crielaard W, Brandt BW, Ramage G. The oral microbiome of denture wearers is influenced by levels of natural dentition. PLos One. 2015; 10:e0137717. doi:10.1371/journal.pone.0137717. PMID: 26368937

94. Ximenez-Fyvie LA, Haffajee AD, Socransky SS. Comparison of the microbiota of supra- and subgingival plaque in subjects in health and periodontitis. J Clin Periodontol. 2000; 27:648–657. PMID: 10983598

95. Ximenez-Fyvie LA, Haffajee AD, Socransky SS. Microbial composition of supra- and subgingival plaque in subjects with adult periodontitis. J Clin Periodontol. 2000; 27:722–732. PMID: 11034118

96. Darout IA, Albandar JM, Skaug N, Ali RW. Salivary microbiota levels in relation to periodontal status, experience of caries and miswak use in Sudanese adults. J Clin Periodontol; 2002; 29:411–420. PMID: 12060423

97. Bostrom L, Bergstrom J, Dahlen G, Linder LE. Smoking and subgingival microflora in periodontal disease. J Clin Periodontol; 2001; 28:212–219. PMID: 11284533

98. Haffajee AD, Socransky SS. Relationship of cigarette smoking to the subgingival microbiota. J Clin Periodontol 2001; 28:377–388. PMID: 11350499

99. Haffajee AD, Bogren A, Hasturk H, Feres M, Lopez NJ, Socransky SS. Subgingival microbiota of chronic periodontitis subjects from different geographic locations. J Clin Periodontol; 2004; 31:996–1002. PMID: 15491316

100. Craig RG, Boylan R, Yip J, Bamgboye P, Koutsoukos J, Mijares D, Ferrer J, Imam M, Socransky SS, Haffajee AD. Prevalence and risk indicators for destructive periodontal diseases in 3 urban American minority populations. J Clin Periodontol. 2001; 28: 524–535. PMID: 11350519

101. Feres M, Haffajee AD, Allard K, Som S, Socransky SS. Change in subgingival microbial profiles in adult periodontitis subjects receiving either systemically-administered amoxicillin or metronidazole. J Clin Periodontol. 2001; 28:597–609. PMID: 11422580

102. Sakellari D, Belibasakis G, Chadjipadelis T, Arapostathis K, Konstantinidis A. Supragingival and subgingival microbiota of adult patients with Down’s syndrome. Changes after periodontal treatment. Oral Microbiol Immunol. 2001; 16:376–382. PMID: 11737662

103. Socransky SS, Haffajee AD. Periodontal microbial ecology. Periodontol 2000. 2005; 38:135–87.

104. Socransky SS, Smith C, Martin L, Paster BJ, Dewhirst FE, Levin AE. Checkerboard DNA–DNA hybridization. Biotechniques. 1994; 17:788–792. PMID: 15853940

105. Pozhitkov AE, Leroux BG, Randolph TW, Beikler T, Flemmig TF, Noble PA. Towards microbiome transplant as a therapy for periodontitis: an exploratory study of periodontitis microbial signature contrasted by oral health, caries and edentulism. BMC Oral Health. 2015; 15:125. doi:10.1186/s12903-015-0109-4. PMID: 26468081

